# Co-designing sequence and structure of functional *de novo* enzymes with EnzyGen2

**DOI:** 10.64898/2026.03.02.709205

**Authors:** Zhenqiao Song, Huichong Liu, Yunlong Zhao, Yang Yang, Lei Li

## Abstract

Proteins underpin essential biological functions across all kingdoms of life. The capacity to design novel proteins with tailored activities holds transformative potential for biotechnology, medicine, and sustainability. However, since protein functions, particularly enzymatic activities, depend on precise interactions with small-molecule ligands, accurately modeling these interactions remains a formidable challenge in *de novo* protein design. Here, we introduce EnzyGen2, a protein foundation model designed for the simultaneous co-design of sequence and structure under ligand-guided functional targeting. Comprising 730 million parameters, EnzyGen2 is trained on 720,993 protein-ligand pairs using multi-task learning objectives that encompass joint prediction of sequence, structure, and protein-ligand interactions. In rigorous *in silico* benchmarks, EnzyGen2 consistently outperforms state-of-the-art baselines, including Inpainting, RFdiffusion/ProteinMPNN, RFdiffusion2/LigandMPNN, and RFdiffusion3/LigandMPNN, as measured by the enzyme-substrate prediction score, AlphaFold2 confidence metric, and structural fidelity, while it generates samples 400× faster than prior methods. We further experimentally validated EnzyGen2 across multiple enzyme families, including chloramphenicol acetyltransferase, aminoglycoside adenylyltransferase, and thiopurine *S*-methyltransferase. *De novo* enzymes generated by our family-specific EnzyGen2 exhibited catalytic activities comparable to or exceeding those of natural enzymes, while retaining substantial novelty with sequence identities as low as 51.6%. These results establish EnzyGen2 as a robust Artificial Intelligence-based tool for functional enzyme design, demonstrating the power of large protein foundation models to create high-performance, novel biocatalysts.

## 1 Introduction

Proteins are vital biomacromolecules of life, executing complex biological functions. A fundamental frontier in protein biochemistry is the *de novo* design of novel proteins with bespoke functions, particularly for catalytic and therapeutic applications [1, 2]. Central to these functions are the interactions between proteins and small-molecule ligands, which underpin the majority of biochemical signaling and metabolism [3–6]. A broad spectrum of proteins, particularly enzymes, capitalize on precise and dynamic protein-ligand interactions to achieve remarkable catalytic efficiency and selectivity in biochemical functions. Consequently, designing proteins capable of binding small-molecule ligands with high affinity and specificity is a primary objective of protein engineering. For enzymes acting on small-molecule substrates, productive catalysis depends on accurate substrate binding, which governs both reaction specificity and rate. Furthermore, cofactor-dependent enzymes require effective binding of their small-molecule cofactors to facilitate chemically demanding transformations. This necessity to capture and optimize protein-ligand interactions adds a critical layer of complexity, rendering the generation of functional *de novo* enzymes a formidable challenge.

Recent advances in deep learning methods have provided powerful generative and discriminative tools for biomolecular design [7–15]. To date, most experimentally validated *de novo* protein design strategies follow a two-stage paradigm [16–18]: protein backbones are first generated to meet structural objectives [19–23], followed by amino-acid sequence design to stabilize the target folding [24–32]. While successful, this sequential approach often fails to capture the reciprocal dependencies between protein sequence and structure that drive biological functions. Although emergent joint sequence-structure co-design approaches have begun to address this limitation [33–36], most existing models lack explicit ligand-binding constraints. The development of a general, ligand-aware *de novo* protein design model capable of simultaneously optimizing sequence and structure across diverse protein families remains a critical and largely unmet need.

Here, we introduce EnzyGen2, a 730-million-parameter foundation model for the simultaneous design of protein sequences and structures under ligand-guided functional targeting. EnzyGen2 employs an interleaved neural network architecture, combining Transformer layers [37] to capture long-range sequence dependencies, equivariant graph neural network layers [38] to model three-dimensional (3D) structural geometries, and a protein-ligand interaction module to enforce the ligand-binding specificity. By leveraging heterogeneous inputs, including small-molecule ligands, functionally important residues, and taxonomic identifiers, EnzyGen2 steers the design process toward evolutionarily plausible sequence spaces, substantially narrowing the search space for functional candidates.

A significant barrier to developing ligand-aware models has been the scarcity of high-quality complex data, with only ∼20,000 publicly accessible protein-ligand complexes [39]. To overcome this limitation, we constructed an expanded dataset to enable effective training of EnzyGen2. We curated proteins with experimentally validated functions from the Protein Data Bank (PDB) [40] and Swiss-Prot [41], and retrieved their corresponding small-molecule ligands from UniProtKB [42]. This resulted in a dataset of 720,993 protein-ligand pairs, exceeding existing complex datasets by more than an order of magnitude. The vast data scale substantially enhanced EnzyGen2’s performance in both *in silico* benchmarks and experimental assays. In *in silico* evaluations, including enzyme-substrate interaction scores [43], AlphaFold2 confidence (pLDDT) [7], and structural fidelity (fraction of designs with root-mean-square deviation (RMSD) *<*2Å), EnzyGen2 consistently outperforms strong, established baselines such as In-painting [33], RFdiffusion/ProteinMPNN [23, 29], RFdiffusion2/LigandMPNN [6, 44], and RFdiffusion3/LigandMPNN [6, 45], while it generates samples 400× faster than prior methods.

Finally, we demonstrate the generality of EnzyGen2 through the experimental validation across three structurally and functionally diverse enzyme families, each operating through distinct catalytic mechanisms and substrate specificities: chloramphenicol acetyltransferase (CAT), aminoglycoside adenyltransferase (AadA), and thiopurine S-methyltransferase (TPMT). Using a rigorous candidate selection pipeline, we generated *de novo* enzymes with catalytic efficiencies comparable to or superior than those of their natural counterparts, despite sharing low sequence identities of 52.4%, 51.6%, and 58.5% with the closest natural enzymes for CAT, AadA, and TPMT, respectively. To our knowledge, this work represents the first experimental demonstration of generative artificial intelligence models for the *de novo* design of these enzyme classes, highlighting the transformative potential of generative protein design in creating functional *de novo* enzymes with activities rivaling natural enzymes. More broadly, this work establishes a new tool for function-driven, ligand-guided protein sequence-structure co-design, opening the door to creating novel enzymes beyond those accessible through natural evolution.

## 2 Results

### 2.1 Overview of EnzyGen2, a Ligand-aware Protein Sequence and Structure Co-design Model

EnzyGen2 is a protein foundation model capable of co-designing sequence and 3D structure given a target binding ligand. The model interleaves Transformer layers and *k*-nearest neighbor equivariant graph neural network layers, the former to capture global amino acid correlations and the latter to model structural geometries (Fig. 1(a)). To steer the candidate function, EnzyGen2 leverages heterogeneous data inputs, including: *1*) automatically identified functionally important residues, which provide precise guidance for specific functions without requiring a *priori* expert annotation; *2*) NCBI taxonomic identifiers (e.g. 562 for *E. coli*), which allow the model to learn organism-specific amino acid combinations, thereby significantly constraining the design search space to evolutionarily plausible regions; and *3*) protein-bound ligands, which are incorporated to capture the fundamental physics and geometry of protein-ligand interactions. EnzyGen2 was trained using a multi-task learning framework [46] comprising masked protein sequence prediction (**ℒ**_seq_), masked backbone structure reconstruction (**ℒ**_str_) and protein-ligand interaction prediction (**ℒ**_bind_). These objectives are jointly optimized through a weighted loss function, where the coefficients *α, β*, and *λ* balance the relative contributions of each component:

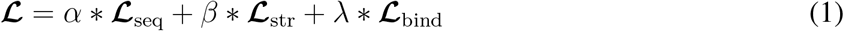

**Fig. 1.**
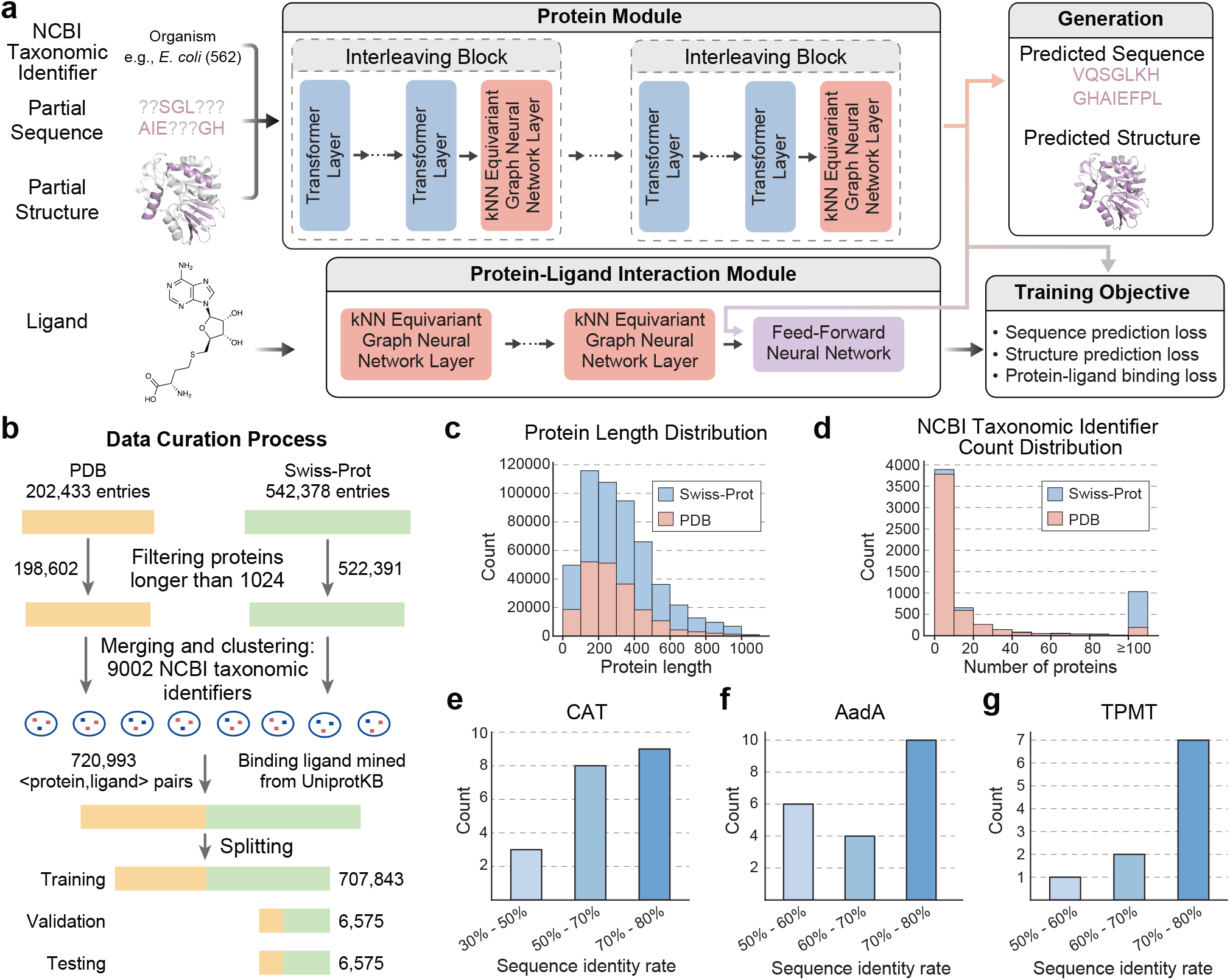
Overview of EnzyGen2 and the curated protein-ligand dataset. **a**, The EnzyGen2 architecture. The model takes an NCBI taxonomic identifier, a partial sequence and 3D structure for functionally important residues, and a ligand as the input. It generates the entire protein sequence and backbone structure. EnzyGen2 includes two modules, a protein module for protein sequence and backbone structure modeling and a ligand module for molecular interaction. The overall model is trained with a set of three losses: a sequence prediction loss, a structure prediction loss and a protein-ligand binding loss. **b**, The training data curation process. **c**, Distribution of protein lengths in our training data: both PDB and Swiss-Prot subsets show consistent distributions, with the majority of proteins ranging from 200 to 600 residues. **d**, Size distributions according to NCBI taxonomic identifiers. Most categories contain fewer than 20 proteins, with a long tail extending 100. **e-g**, Sequence identity rate distributions of *de novo* designed candidates relative to natural homologs within the same enzyme families.

To pretrain EnzyGen2, we curated a comprehensive dataset by merging experimentally validated protein sequences from the PDB and Swiss-Prot (Fig. 1(b)). Backbone structures for PDB entries were obtained from crystal structures, while Swiss-Prot proteins utilized predicted folds from the AlphaFold Protein Structure Database [47]. After filtering proteins longer than 1,024 amino acids, the final dataset comprised 720,993 unique proteins. As illustrated in Fig. 1(c), the majority of proteins in our dataset ranged between 200 and 600 amino acids in length. Ligand information was extracted from UniProtKB [48]. Proteins with a documented ligand were assigned a positive binding label, while those without a documented ligand were assigned a negative label by randomly sampling a small molecule. To steer the model with evolutionary taxonomy, proteins were clustered by their NCBI taxonomic identifiers, yielding 9,002 distinct clusters (Fig. 1(d)) with 263 clusters containing more than 500 proteins. From these large clusters, 25 proteins were randomly sampled for the validation and test sets, resulting in a final dataset partition of 707,843 protein-ligand pairs for training and 6,575 pairs each for validation and testing.

To adapt our pretrained model to specific protein families with minimal computational overhead, we fine-tuned EnzyGen2 on family-specific datasets. In this study, we focused on the *de novo* design of enzymes. We curated datasets comprising 3,121 entries of chloramphenicol acetyltransferase (CAT), 2,499 entries of aminoglycoside adenylyltransferase (AadA), and 6,770 entries of thiopurine *S*-methyltransferase (TPMT) from the Rhea reaction database [49]. Each dataset was randomly split into training and validation sets with a 9:1 ratio. Fine-tuning was optimized only on the masked sequence prediction loss (**ℒ**_seq_) and masked structure reconstruction loss (**ℒ**_str_), as the substrates for these enzyme families remained the same within each enzyme family.

### 2.2 Systematic Selection of Diverse, Novel, Functional Enzyme Candidates

To identify high-quality candidates for experimental validation, we implemented a multi-stage screening pipeline integrating structural, functional metrics, and evolutionary metrics (Supplementary Fig. S1). The workflow ranks candidates using function scores, structural stability, sequence log-likelihood (representing evolutionary plausibility), diversity and novelty. The functional assessment combined the enzyme-substrate prediction (ESP) score [43] with an adversarial discriminator score [50]. The latter was designed to distinguish natural enzymes from EnzyGen2-generated ones. A higher discriminator score indicates that the generated enzyme closely resembles natural enzymes in function, although not necessarily in sequence identity. To train this adversarial discriminator, we generated a set of artificial enzymes using nucleus sampling [51] with *p* ∈ {0.2, 0.4, 0.6, 0.8}, labeled natural enzymes as 1, artificial enzymes as 0, and fine-tuned a 650M-parameter ESM2 model [14] for binary classification.

The structural stability of the designed enzymes was estimated using pLDDT scores from ESM-Fold [14]. The evolutionary scores of sequences was quantified by their log-likelihood within EnzyGen2. Diversity and novelty scores were calculated as the minimum sequence difference ratio (1 - maximum sequence identity rate after pairwise alignment) relative to previously selected candidates and natural enzymes within the same family, respectively.

Candidates were selected through a stringent filtering pipeline starting from an initial generated pool. Samples are selected based on the following criteria: ESP score *>* 0.6, adversarial discriminator score *>* 0.6, and pLDDT *>* 80. The surviving sequences were grouped into subsets based on their maximum sequence identity to natural enzymes, categorized into bins of 30-40%, 40-50%, 50-60%, 60-70%, and 70-80%. Candidates with maximum identity rate below 30% or above 80% were excluded to ensure samples were neither excessively divergent from known folds nor overly similar to natural enzymes. Within each bin, sequences were ranked by their log-likelihood scores. The top-ranked candidates were further examined by experts based on the geometric alignment of their active sites with natural templates, while ensuring a minimum sequence diversity score of 10% among all previously selected candidates.

Following this workflow, we selected 20 chloramphenicol acetyltransferase (CAT), 20 aminoglycoside adenylyltransferase (AadA), and 10 thiopurine *S*-methyltransferase (TPMT) candidates for wet-lab testing. The maximum sequence identity rate of these candidates to natural homologs within the same family ranged from 30% to 80% (Fig. 1(e-g)), demonstrating EnzyGen2’s capacity to traverse diverse regions of the sequence space while achieving high functionality.

### 2.3 *In Silico* Evaluation of the Pretrained EnzyGen2

To benchmark the performance of our pretrained model, we conducted a comparative analysis against the state-of-the-art protein sequence and structure co-design methods. Focusing on the ten most frequent enzyme families in our test set, we first compared EnzyGen2 to Inpainting [33], providing both models with the same functionally important residues as input to ensure a rigorous comparison. As illustrated in Fig. 2(a), our EnzyGen2 achieved superior enzyme-substrate prediction (ESP) scores in 7 of 10 categories and a higher proportion of designs with RMSD *<* 2Å in 9 out of 10 categories. This enhanced functional performance stems from the explicit protein-ligand interaction loss (**ℒ**_bind_) during the training of EnzyGen2. Furthermore, EnzyGen2 designs yielded higher AlphaFold2 pLDDT scores [7] in 9 of the 10 categories, confirming superior structural stability. We further assessed self-consistency (scRMSD)— a key metric for design viability — by measuring the RMSD between the designed structures and AlphaFold2 predictions of the designed sequences (Fig. S2). EnzyGen2 exhibited lower scRMSD values in 8 of the 10 enzyme families, achieving *<* 2Å agreement for 4 families, which highlights a substantial higher level of sequence-structure coherence than Inpainting.

**Fig. 2.**
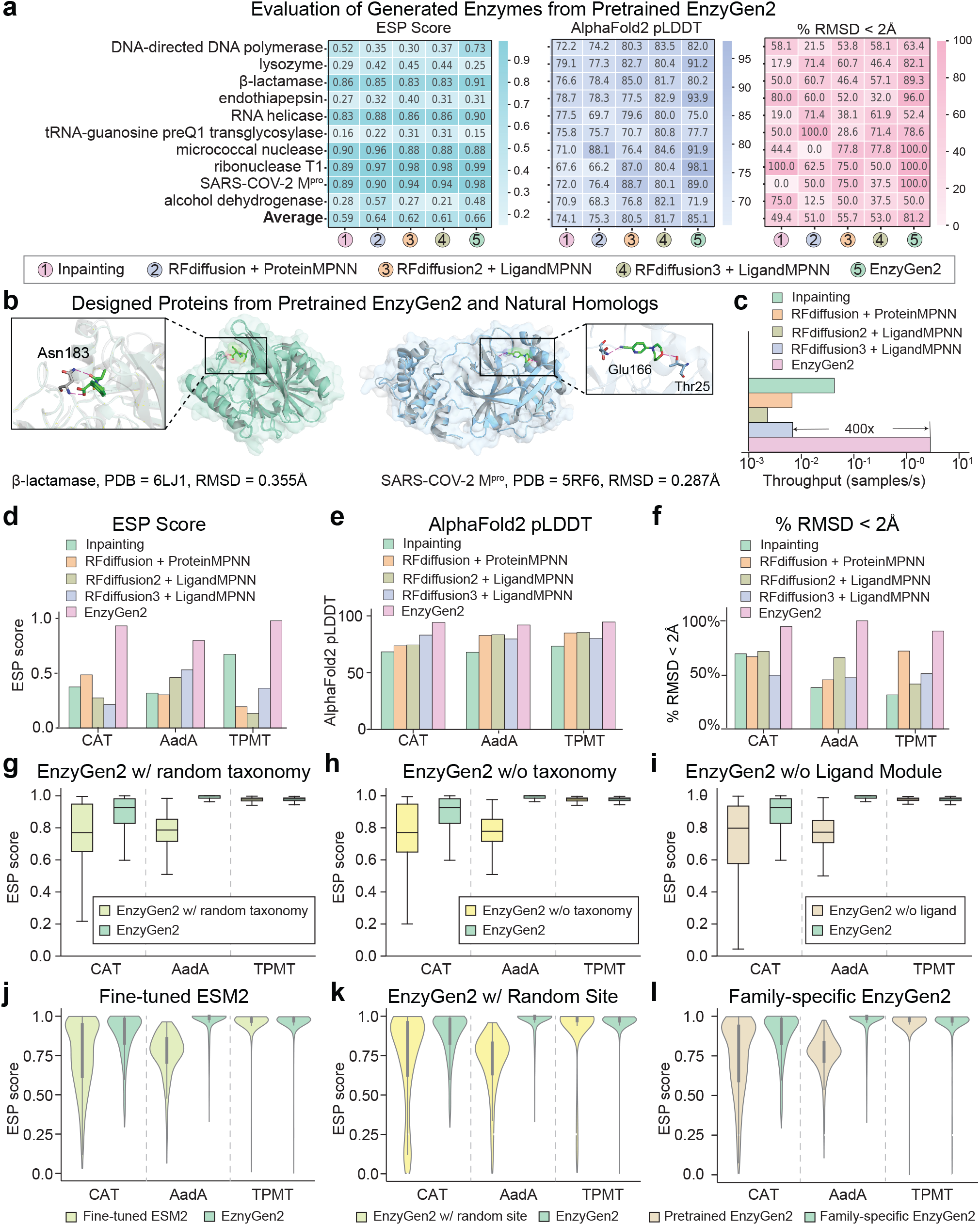
*In silico* evaluation results and ablation analysis of EnzyGen2. **a**, The pretrained EnzyGen2 outperforms state-of-the-art baselines on all three averaged metrics across 10 frequent enzyme families. **b**, Structural superposition of *de novo β*-lactamase and SARS-COV-2 Mpro designs (colored) against natural homologs (grey). **c**, Throughput (log10 scale) of all models. **d-f**, Performance of family-specific EnzyGen2 models for CAT, AadA, and TPMT across ESP score, AlphaFold2 pLDDT, and fraction of RMSD < 2Å against natural homologs. **g-l**, Ablation analysis identifying the impact of NCBI taxonomic identifiers, ligand-binding module, sequence structure co-design framework, functionally important residue constraint, and family-specific fine-tuning.

We next compared our method against the widely used RFdiffusion/ProteinMPNN pipeline [17, 23, 29, 52]. We provided the main-chain backbone coordinates of functionally important residues to RFdiffusion, followed by sequence generation via ProteinMPNN. As shown in Fig. 2(a), EnzyGen2 out-performed this pipeline in higher enzyme-substrate prediction (ESP) scores (5 of 10 categories) and structural fidelity (RMSD *<* 2Å in 8 of 10 categories). We noticed that the RFdiffusion/ProteinMPNN pipeline failed to generate any proteins with an RMSD *<* 2Å for micrococcal nuclease. To investigate this failure mode, we calculated the binding affinity using Gnina [53]. The pipeline yielded a poor average binding energy of −2.18 kcal/mol compared to other models, suggesting a distinct limitation of the pipeline for this specific enzyme class. Across all categories, EnzyGen2 produced designs with an mean AlphaFold2 pLDDT score of 85.10, significantly exceeding the standard high-stability threshold of 80 [54], thus confirming its ability to design *de novo* stable, foldable enzymes.

We further benchmarked EnzyGen2 against the RFdiffusion2/LigandMPNN pipeline [6, 44], representing the state of the art in enzyme design. To generate the structural backbones, we first provided RFdiffusion2 with the target ligand and functionally important residues, sourcing these initial coordinates from either experimentally validated PDB complexes or AlphaFold3 predictions [15]. These generated structures were subsequently used as inputs for LigandMPNN to design the final sequences. Even against this strong baseline, EnzyGen2 demonstrated superior average performance across all metrics (Fig. 2(a)), including ESP scores, AlphaFold2 pLDDT, and the fraction of high-fidelity designs (RMSD *<* 2Å). Collectively, these results establish EnzyGen2 as a robust platform for functional *de novo* design, capable of accurately coordinating enzymes with substrate and cofactor binding.

Finally, we evaluated EnzyGen2 against the RFdiffusion3/LigandMPNN pipeline [6, 45], which serves as a benchmark for *de novo* protein design under ligand-binding constraints. Following the standard protocol for this pipeline, ligand coordinates and functionally important residues were provided to RFdiffusion3 for structural generation, after which LigandMPNN was employed to derive sequences compatible with the designed scaffolds. Despite the competitive performance of this state-of-the-art baseline, EnzyGen2 demonstrated superior performance across all evaluation metrics (Fig. 2(a)). Notably, EnzyGen2 achieved higher ESP scores in 8 of 10 categories, enhanced structural stability in 5 categories, and superior structural fidelity (RMSD *<* 2Å) in 9 of 10 categories. These findings validate EnzyGen2 is a versatile platform for the *de novo* design of diverse protein families.

To further evaluate the computational efficiency of EnzyGen2, we measured the throughput (samples/s) across the ten most frequent enzyme families (Fig. 2(c)). EnzyGen2 demonstrated a substantial speed advantage, achieving speedups of 72×, 482×, 1,420×, and 469× compared to Inpainting, RFdiffusion/ProteinMPNN, RFdiffusion2/LigandMPNN, and RFdiffusion3/LigandMPNN, respectively. These results underscore a highly accelerated generation process, highlighting the practical utility of EnzyGen2 for the rapid *de novo* design of enzymes.

To better visualize EnzyGen2’s design capabilites, we highlight two case studies (Fig. 2(b)). We first employed AlphaFold2 to predict the structure of the designed proteins. This predicted structure was then aligned to the reference protein structure from the PDB with the corresponding ligand using PyMOL [55]. As shown in Fig. 2(b), both designs achieved high structural alignment (RMSD *<* 0.5Å) and successfully exhibited essential hydrogen bonds with the corresponding ligands (in purple). These results provide strong evidence that EnzyGen2 can generate novel enzymes that adopt targeted functional folds while maintaining the precise physicochemical relations required for catalysis.

### 2.4 *In Silico* Validation of Family-specific Enzyme Designs

To demonstrate the adaptable design capabilities of EnzyGen2 for family-specific enzyme design, we fine-tuned the pretrained EnzyGen2 on three representative enzymes: chloramphenicol acetyltransferase (CAT), aminoglycoside adenylyltransferase (AadA), and thiopurine *S*-methyltransferase (TPMT). The curation of these fine-tuning datasets is detailed in Section 4.4. For each enzyme family, we mined the functionally important residues of all natural enzymes. We then used the nucleus sample method [51] to generate candidates from EnzyGen2 (the probability threshold *p* ∈ {0.2, 0.4, 0.6, 0.8}) and identified qualified candidate sets for each family using the systematic selection pipeline described in Section 2.2. We evaluated all baseline methods against the same natural enzyme reference sets for fair comparison.

As illustrated in Fig. 2(d-f), the family-specific EnzyGen2 consistently achieved superior performance across all computational metrics. Specifically, our model outperformed all baselines in enzyme-substrate prediction (ESP) scores, AlphaFold2 pLDDT, and structural fidelity (fraction of designs with RMSD *<* 2Å). Notably, the fine-tuned models achieved an average ESP score exceeding 0.6, an AlphaFold2 pLDDT greater than 90, and the proportion of RMSD *<* 2Å designs surpassing 90% across all enzyme families. These results provide robust *in silico* evidence that fine-tuned EnzyGen2 models can design *de novo* enzymes with high structural stability and a strong likelihood of functional success in subsequent experimental validation.

### 2.5 Ablation Studies Identify Key Drivers of Functional Design Performance

To systematically evaluate the contribution of each component in EnzyGen2, we conducted a series of ablation experiments. We isolated the effects of the NCBI taxonomic identifier, the protein-ligand interaction prediction module, the sequence-structure co-design framework, functionally important residue constraints, and the fine-tuning strategy.

To assess the functional role of the NCBI taxonomic identifier in constraining the design search space, we performed a two-step evaluation. First, we tested taxonomic specificity by randomly assigning NCBI identifiers to design tasks and examining the resulting enzyme-substrate prediction scores (ESP) across three enzyme families (Fig. 2(g)). Candidates were generated using nucleus sampling with a probability threshold of *p* = 0.4. The observed consistent decrease in ESP scores suggests that each NCBI identifier corresponds to a specific, evolutionarily constrained set of functionally important residues; randomizing this input effectively disrupts this essential correspondence. Second, we assessed the identifier’s overall necessity by removing the NCBI identifiers entirely during candidate generation (Fig. 2(h)). In this setting, the enzyme-substrate interaction (ESP) scores decreased across all enzyme families. Together, these findings confirm that the NCBI taxonomic identifiers are essential for reducing the combinatorial search cost and providing vital evolutionary guidance.

To further evaluate the utility of ligand information, we trained a variant of our model that excluded the protein-ligand interaction module and the associated binding prediction loss **ℒ**_bind_. Candidates were generated using nucleus sampling with a probability threshold of *p* = 0.4. The resulting distributions of ESP scores are depicted in Fig. 2(i). We observed a substantial decrease in small-molecule ligand interaction scores. This performance drop unequivocally demonstrates the irreplaceable role of the proteinligand interaction constraint in steering our model toward successful ligand-binding protein design.

To rigorously assess the synergy between sequence and structural modeling, we compare the full EnzyGen2 architecture and two ablated variants. The first variant removed the Transformer layers and protein-ligand interaction module, relying exclusively on *k*-nearest neighbor equivariant graph neural networks (*k*NN-EGNN); the second eliminated the *k*NN equivariant graph neural network layers and protein-ligand interaction module, relying solely on protein sequences (i.e., Fine-tuned ESM2). Consistent with previous reports [56], the removal of Transformer layers led to unstable training dynamics, particularly for longer proteins. Consequently, we focused our analysis on the sequence-only variant (Fine-tuned ESM2, Fig. 2(j)). This model exhibited uniformly lower ESP scores compared to the full codesign model, which decisively demonstrates that the integration of 3D backbone structure information is crucial for generating proteins with high-fidelity ligand-binding functions.

To assess the utility of functionally important residues, we conducted an ablation study where we replaced the functionally important residues with randomly selected residues of equivalent size during candidate generation. Candidates were generated using nucleus sampling with a probability threshold of *p* = 0.4. The resulting ESP scores (Fig. 2(k)) showed a consistent and significant performance drop across all enzyme families. Critically, this degradation was the most pronounced observed when compared to the removal of any other module (Fig. 2(h-j)). These findings decisively demonstrate that functionally important residues provide the primary, indispensable guidance for *de novo* protein design, effectively dictating the attainment of expected target functions.

We have demonstrated the generalization ability of our pretrained model across diverse enzyme families (Fig. 2(a)). To investigate whether additional fine-tuning improves design quality, we evaluated and compared the ESP scores of both the pretrained and fine-tuned models on three enzyme families (Fig. 2(l)). While the pretrained EnzyGen2 already showed good performance, with ESP scores consistently above 0.6 (indicating positive binding) for all families, further fine-tuning significantly enhanced the design quality. The fine-tuned model achieved scores above 0.8 across all families, confirming that family-specific fine-tuning is a powerful tool for maximizing the success rate of *de novo* enzyme design.

### 2.6 Experimental Validation and Biochemical Characterization of Three Classes of *De Novo* Designed Enzymes

To evaluate the catalytic activity of EnzyGen2-designed enzymes, we selected two antibiotic resistance enzymes, including chloramphenicol acetyltransferase (CAT; Fig. 3(a)) and aminoglycoside adenylyl-transferase (AadA; Fig. 3(b)). CAT is an acyltransferase capable of acetylating the hydroxyl group of chloramphenicol for its conversion [57]. On the other hand, AadA is an adenylyltransferase that modifies aminoglycoside by transferring an adenynyl group (AMP) from adenosine triphosphate (ATP) to a hydroxy group of the aminoglycoside substrate [58]. Genes encoding the *de novo* designed CAT and AadA were synthesized and inserted into the pET-28a(+) and pET-22b(+) vectors respectively between the NcoI and XhoI restriction sites and introduced into *E. coli* by electroporation. Transformants were selected on LB agar plates containing the appropriate antibiotics and lactose. Colony formation on selective media served as evidence of successful transformation and functional expression of the EnzyGen2-designed antibiotic resistance enzymes. Seven of the twenty designed CATs enabled *E. coli* survival on chloramphenicol-containing agar plates, while eight of the twenty designed AadAs conferred resistance to aminoglycoside antibiotics, collectively confirming the catalytic activity of these *de novo* designed enzymes.

**Fig. 3.**
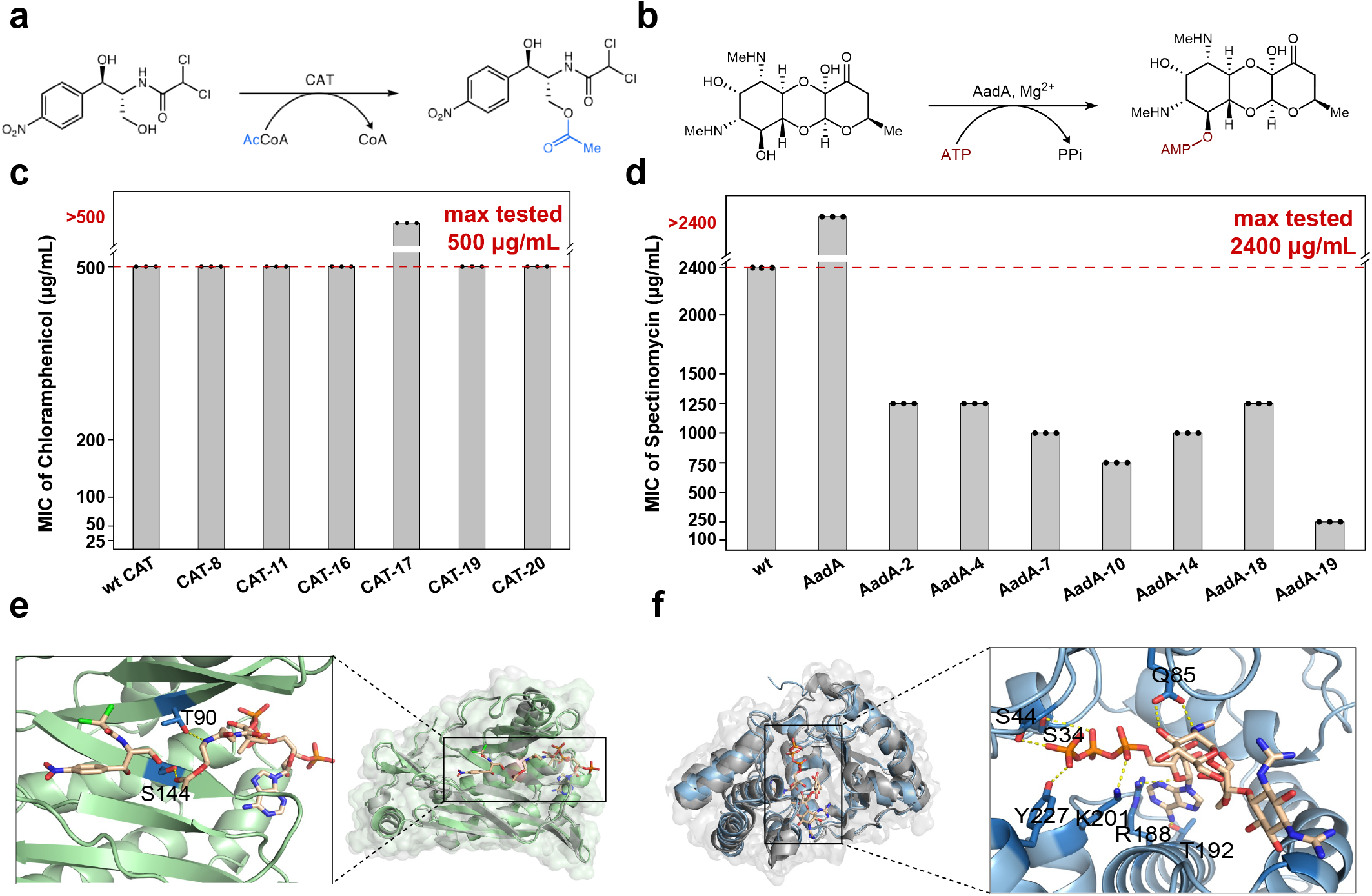
Biochemical and structural validation of *de novo* chloramphenicol acetyltransferase (CAT) and aminoglycoside adenylyltransferase (AadA) enzymes. **a**, Reaction mechanism for CAT-catalyzed acetylation of chloramphenicol using acetyl-CoA as the acyl donor. **b**, Reaction mechanism for AadA-catalyzed adenylylation of antibiotics using ATP as the adenylyl donor. **c-d**, Minimum inhibitory concentration (MIC) determination for *E. coli* strains expressing (**c**) *de novo* CAT variants and (**d**) *de novo* AadA variants. All experiments were conducted in biological triplicates. **e-f**, Structural superposition of (**e**) the designed CAT-17 (green) and (**f**) the designed AadA-2 (blue) against their respective natural homologs (grey; PDB 6X7Q and 6FZB).

Next, we determined the minimum inhibitory concentrations (MICs) of *E. coli* strains harboring individual designed antibiotic degrading genes to assess their relative activities. All active *de novo* CATs exhibited resistance levels comparable to the wild-type *E. coli* CAT. Notably, *E. coli* harboring *de novo* CAT-17 grew robustly in LB medium containing 500 *µ*g/mL chloramphenicol, a concentration at which the strain expressing wild-type *Ec*CAT failed to survive (Fig. 3(c)). Among the *de novo* designed AadA enzymes, AadA-2 allowed *E. coli* growth in LB medium supplemented with up to 2400 *µ*g/mL spectinomycin, representing a substantial enhancement in resistance relative to the wild-type AadA from *Salmonella typhimurium* (Fig. 3(d)).

To further validate the enzymatic activity of the designed proteins, we examined the acylative inactivation of chloramphenicol using our designed CATs under whole-cell conditions. Bacterial metabolites were then analyzed by liquid chromatography-mass spectrometry (LC-MS). All seven designed CATs that displayed substantial activity in our *E. coli* survival assays produced acetylated chloramphenicol, directly confirming the acetyl transferase activity of these *de novo* CATs (Fig. S3-S9). Finally, structural superposition of our designed enzymes onto their natural homologs, CAT-17 against *Ec*CAT from *E. coli* (PDB: 6X7Q) and AadA-2 against *Se*AadA from *Salmonella enterica* (PDB: 6FZB), yielded RMSD values of 0.54 Å and 1.08 Å, respectively (Fig. 3(e-f)). These low deviations confirm that the EnzyGen2-designed enzymes preserve the active-site geometry required for catalytic function.

To further demonstrate the utility of EnzyGen2 for the design of other biotechnologically useful enzymes, we focused our efforts on thiopurine *S*-methyltransferase (TPMT) [60], a class of enzymes with excellent potential for the regeneration of *S*-adenosylmethionine (SAM) [59, 61, 62], a widely used cofactor in biotransformations [63–66]. Although the native function of TPMT is to transfer the sulfonium methyl group from SAM to purine substrates [67], these enzymes also demonstrated substantial halogen methyltransferase (HMT) activity in previous studies [62, 68]. In particular, TPMTs can catalyze methyl transfer of methyl from methyl halide, most notably methyl iodide as well as methyl sulfonates, to *S*-adenosylhomocysteine (SAH) (Fig. 4(a)). This reaction enables the *in vitro* regeneration of SAM, a ubiquitous methyl donor that underpins biocatalytic platforms for regio- and/or enantioselective methylation which are challenging with chemical systems [69–77]. Although several native HMTs capable of methylating SAH using methyl iodide have been identified, their practical application is limited by their low expression level in *E. coli* and variable catalytic efficiency. We therefore envisioned that *de novo* designed TPMTs could serve as efficient SAM-regeneration enzymes and overcome these limitations..

**Fig. 4.**
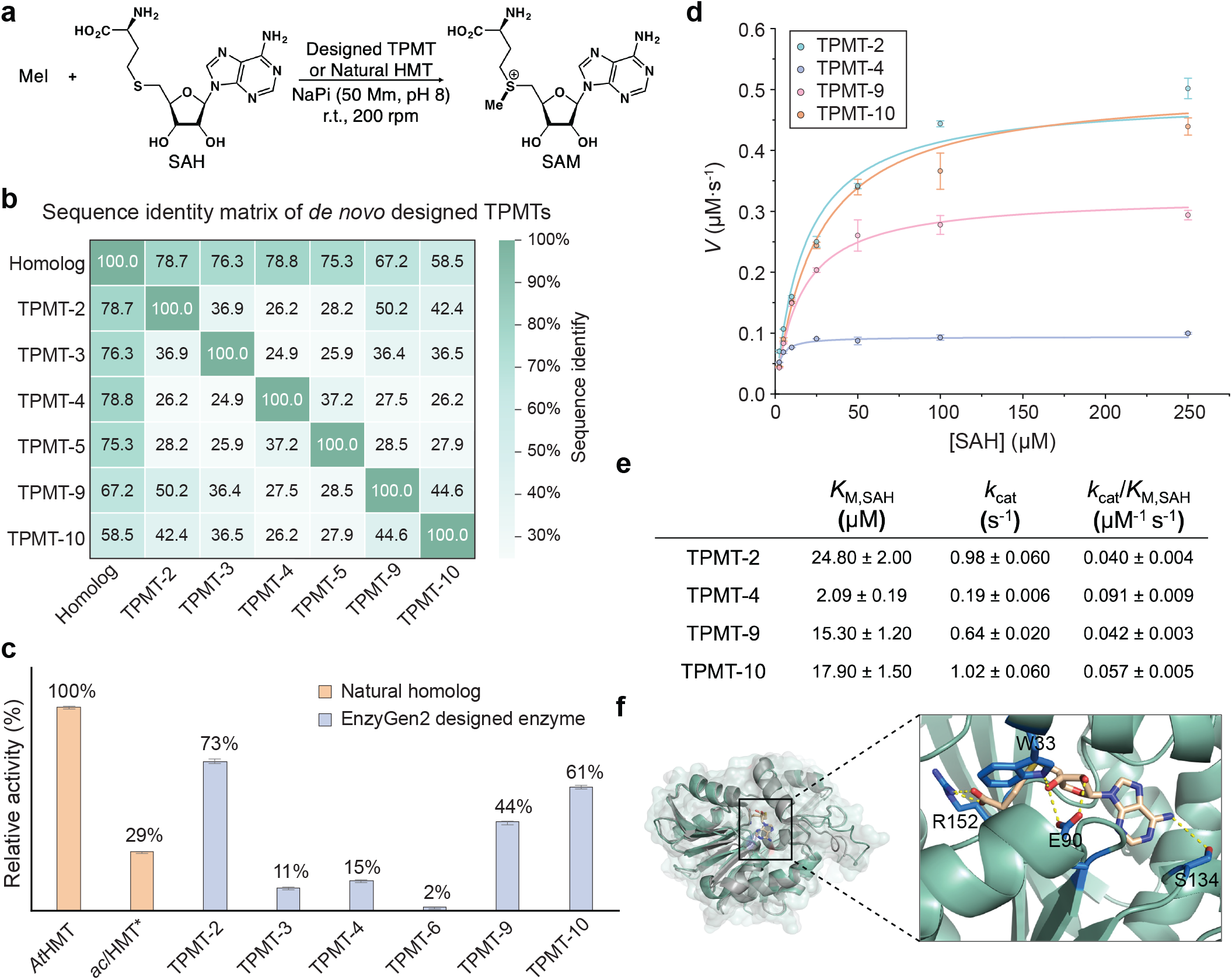
*De novo* design, experimental validation and kinetic characterization of thiopurine *S*-methyltransferases (TPMTs) for SAM-regeneration. **a**, TPMT/HMT-catalyzed regeneration of SAM from SAH and methyl iodide. **b**, Sequence identity matrix of EnzyGen2-designed TPMT variants. **c**, Relative activities of EnzyGen2-designed enzymes. Initial rates were determined by measuring product yields after 1 min under standard reaction conditions. Reaction conditions: 10 mM MeI, 250 *µ*M SAH, 0.5 *µ*M TPMT, 100 mM NaPi (pH = 8.0), 25 °C. The activity data for *acl*HMT were obtained from the literature [59]. **d-e**, Michaelis-Menten kinetics of the four most active *de novo* designed TPMTs. Reactions were performed under standard conditions with varying SAH concentrations. **f**, Structural superposition the designed TPMT-4 (green) against its natural homolog, PDB 2BZG (grey).

To this end, we used EnzyGen2 to generate 10 *de novo* TPMTs exhibiting 58.4% – 78.8% sequence identity with their closest natural homologs in the reference library (Fig. 4(b)). The catalytic activities of these *de novo* TPMTs were subsequently experimentally evaluated in the SAM-regeneration reaction using SAH and methyl iodide (MeI). Similar to our work with CATs and AadAs, *de novo* designed TPMTs were cloned into the pET-28a(+) vector between the NcoI and XhoI restriction sites and overexpressed in *E. coli* BL21(DE3) strain. Expression levels of these *de novo* TPMTs were assessed by sodium dodecyl sulfatepolyacrylamide gel electrophoresis (SDS-PAGE) analysis, revealing that six out of the ten designed TPMTs were expressed at a high level in *E. coli* (Fig. S10). These TPMTs were subsequently characterized using a series of enzyme kinetic assays.

Initial reaction rates were measured and normalized to that of the halomethyltransferase from *Ara-bidopsis thaliana* (*At*HMT), which was set to 1. The *de novo* TPMTs exhibited relative activities ranging from 2% to 73%. To provide broader benchmarking, we additionally included *acl*HMT, a halomethyl-transferase (HMT) from *Aspergillus clavatus* known for its broad substrate promiscuity and synthetic utility [77]. Among the *de novo* designed TPMTs, including TMPT-2, TMPT-9 and TMPT-10, higher activity than the widely used natural enzyme *acl*HMT (Fig. 4(c)), demonstrating their potential for application in biocatalysis.

Further Michaelis-Menten kineticcharacterization was carried out with the four top-performing TPMTs, including TPMT-2, 4, 9 and 10 (Fig. 4(d-e))). TPMT-4 exhibited a *K*_M, SAH_ value of (2.09±0.19) *µ*M and a *k*_cat_ value of (0.19±0.006) *s*^−1^, corresponding to a *k*_cat_/*K*_M_ value of 0.091±0.009 *µ*M^−1^*s*^−1^, comparable to that of *Ct*HMT (*k*_cat_/*K*_M_ = 0.29±0.09 *µ*M^−1^*s*^−1^), a previously characterized natural HMT from *Chloracidobacterium thermophilum* [69]. In addition, TPMT-10 displayed a higher turnover number *k*_cat_ of (1.02 ± 0.06 *s*^−1^) than that of *Ct*HMT (*k*_cat_ = 0.33± 0.03 *s*^−1^) [69]. Notably, the two best-performing designs displayed distinct kinetic profiles: TPMT-4 achieved high catalytic efficiency through strong substrate binding affinity (low *K*_M, SAH_) coupled with moderate turnover, whereas TPMT-10 (along with TPMT-2 and TPMT-9) exhibited lower affinity but substantially higher turnover rates (high *k*_cat_).

To investigate the origins of these functional differences, we analyzed the closest natural homologs of *de novo* designed TPMT-4 and TPMT-10 using BLAST searches in the Uniprot database. TPMT-4 aligns closely with TMPT (Uniprot ID: G3T7V7) and a structurally resolved TPMT (PDB ID: 2BZG), representing a well-characterized enzyme family. Further structural superposition of TPMT-4 onto the crystal structure of a natural TPMT (PDB ID: 2BZG; Fig. 4(f)) provided an RMSD of 0.46 Å, confirming that the designed enzyme adopts the native fold and preserves the geometric features required for catalysis. In contrast, the closest homolog to designed TPMT-10 is an uncharacterized protein (Uniprot ID: A0A1G9ECT0). These two lineages share only ca. 30% sequence identity, indicating that the TPMT family comprises at least two distinct clades, the latter of which remains largely unexplored in biotransformations and biotechnology. The divergence in kinetic parameters observed between TPMT-4 and TPMT-10 correlates well with this phylogenetic separation. Crucially, these results demonstrate that EnzyGen2 can access previously uncharacterized and unexplored enzyme subfamilies and enable exploration of uncharted protein space.

## 3 Discussion

EnzyGen2 is a protein foundation model that accepts multiple flexible combinations of functional intent. By implementing a co-design strategy that jointly models sequence and structure, EnzyGen2 achieves superior conformational consistency, significantly reducing the deviation between the designed backbone and the folded structure of the generated sequence. This high level of structural fidelity suggests that the EnzyGen2 model has captured fundamental biophysical principles governing the sequence-structure-function relationship. Across diverse enzyme families, our pretrained EnzyGen2 consistently outperforms existing state-of-the-art methods. Crucially, the *de novo* enzymes generated by the fine-tuned EnzyGen2 model exhibit catalytic efficiencies comparable to, or exceeding, those of their natural counterparts, despite sharing sequence identities as low as 51.6%. These findings establish EnzyGen2 as a powerful platform for *de novo* enzyme design.

The strong performance of EnzyGen2 can be attributed to its large-scale pretraining on an extensive collection of protein-ligand pairs curated from both the PDB and Swiss-Prot. This broad data coverage endows the model with generalizable design capabilities that extend beyond the specificities of individual enzyme classes. Compared to our previous EnzyGen framework [78], which was primarily tailored for enzyme-specific generation under substrate-binding constraints, EnzyGen2 substantially expands the design landscape to encompass general protein-ligand interaction modeling. This advancement enables the design of functional proteins beyond modeling enzymatic systems and supports a wider spectrum of biochemical interactions involving diverse small-molecule ligands. A key model improvement is the replacement of the enzyme classification (EC) tags used in EnzyGen with hierarchical NCBI taxonomic idenfiers. These NCBI identifiers encode richer evolutionary information across protein families and substantially reduces the effective search space during design. Together with the enlarged pretraining corpus, this architectural and data-level expansion increases the model capacity from 714 million to 733 million parameters, enabling more nuanced representation learning of sequence-structure-function relationships. Both *in silico* benchmarking and experimental validation consistently demonstrate that EnzyGen2 generates high-quality candidates across a broad range of protein families, positioning EnzyGen2 as a versatile foundation model for *de novo* enzyme design.

Despite its strong performance and successful experimental validation, several opportunities remain for further optimization. First, although our protein-ligand interaction constraints are enforced during training and candidate selection, they are not explicitly integrated into the generative process itself. Future work will incorporate iterative optimization strategies, such as simulated annealing [79], guided by functional scoring metrics including enzyme-substrate interaction scores or docking energies. Such approaches would allow candidates to be iteratively optimized for functional scores during the design process. Second, our current pretraining dataset is composed solely of experimentally confirmed data from the PDB and Swiss-Prot. The vast, albeit sparsely annotated, dataset in TrEMBL [41] represents an unexplored resource that could be used through a multi-stage training strategy, in which initial largescale pretraining on TrEMBL is followed by continued training on curated datasets to reinforce functional accuracy. Finally, while EnzyGen2 co-designs amino acid sequences and the *C*_*α*_-based backbone structures, extending the framework to full-atom structural representations could impose more stringent physical constraints and further improve design quality.

In summary, this study introduces EnzyGen2 as a powerful and versatile platform for *de novo* protein design with a demonstrated ability to generate high-quality, functional enzymes. We anticipate that the development of structure-sequence co-design models will further improve design success rates and enable the routine generation of enzymes with high catalytic efficiency and tailored functionality, accelerating the development of biocatalysts for green chemistry, therapeutics, and industrial biotechnology.

## 4 Methods

### 4.1 EnzyGen2 Architecture and Multi-task Learning Objective

Let **𝒜** be the set of 20 common amino acids and **𝒞** the set of all NCBI taxonomic identifiers. A protein of length *N* is represented by its amino acid sequence ***a*** = {*a*_1_, *a*_2_, …, *a*_*N*_ } ∈ **𝒜**^*N*^ and corresponding *C*_*α*_ coordinates ***x*** = [***x***_1_, ***x***_2_, …, ***x***_*N*_ ]^*T*^ ∈ ℝ^*N* ×3^. For each residue *a*_*i*_ ∈ **𝒜**, where *i* ∈ {1, 2, …, *N* }, we denote its one-hot encoding by ***a***_*i*_ = onehot(*a*_*i*_), and define the index set of functionally important residues as **ℱ**. The taxonomic identifier of the protein is represented by *c* ∈ **𝒞**. A ligand is represented as **𝒱** = {*v*_1_, …, *v*_*m*_}, where *m* is the number of its atoms. Each atom is given by *v*_*j*_ = (***h***_*j*_, ***x***_*j*_), where ***h***_*j*_ ∈ ℝ^5^ denotes five pre-computed chemical features such as atom type using RDKit [80] and ***x***_*j*_ ∈ ℝ^3^ its corresponding coordinate, for *j* ∈ {1, 2, …, *m*}. Each protein-ligand pair is associated with a binary label ***y*** indicating whether the ligand binds to the protein. The problem studied in this paper can be formulated as follows: given a taxonomic identifier *c*, a ligand **𝒱**, a functionally important residue index set **ℱ**, its residue set ***a***_**ℱ**_ and the corresponding 3D coordinates ***x***_**ℱ**_, generate a protein sequence ***a*** and *C*_*α*_ coordinates ***x*** of *N* residues satisfying the ligand binding constraint ***y***. Essentially, we aim to learn a generative model *P* (***a, x, y***|***a***_**ℱ**_, ***x***_**ℱ**_, *c*, **𝒱**; ***θ***) with parameters ***θ***:

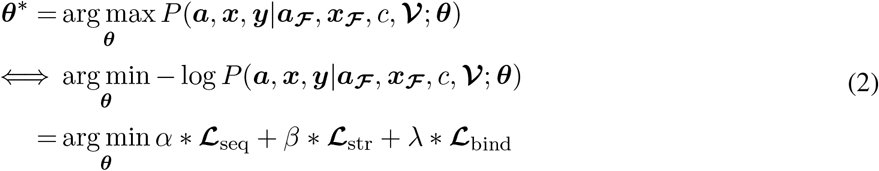

At inference time, we only provide the taxonomic identifier *c*, the functionally important residue types ***a***_**ℱ**_ and coordinates ***x***_**ℱ**_ to simultaneously generate a protein sequence and *C*_*α*_ backbone structure of a desired length.

EnzyGen2 consists of a protein module and a protein-ligand interaction module. The protein module is composed of *L*_*b*_ interleaving blocks to effectively capture the complex dependencies between protein sequences and 3D structures, enabling improved co-design performance. Each interleaving block (Fig. 1(a)) comprises eleven Transformer layers and a *k*-nearest neighbor equivariant graph neural network layer (*k*NN-EGNN). The output of each block serves as the input to the subsequent block, ensuring seamless integration of sequence and structure information. Within each block, the stacked Transformer layers are employed to model the entire sequence dependencies, leveraging a self-attention mechanism to capture contextual relationships effectively:

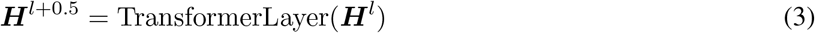

where 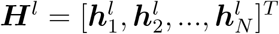 and 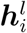 is the *i*^*th*^ residue input at *l*^th^ block. The input residue embeddings for the first layer are either taken from an embedding lookup table ***E*** for functionally important residues, or initialized with a special [mask] token embedding for other residues. The residue embedding will then be enhanced by the NCBI identifier embedding *f* (*c*) corresponding to the desired NCBI taxonomic identifier *c*:

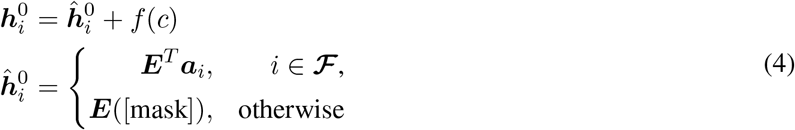

Then a *k*-nearest neighbor equivariant neural network layer (*k*NN-EGNN) in the current block is leveraged to capture the residue correlations in the 3D space:

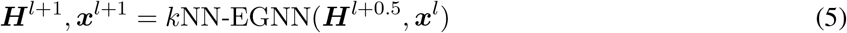

After completing the calculations for all interleaving blocks, the corresponding loss functions for masked sequence prediction (**ℒ**_seq_) and masked structure reconstruction (**ℒ**_str_) can be calculated as follows:

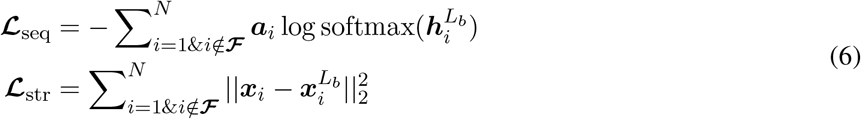

where 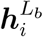 and 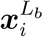 denote the output representation and coordinate of the *i*-th residue from the last interleaving block.

To enforce the ligand-binding specificity constraint during training, we build the protein-ligand interaction module as an equivariant graph neural network consisting of *L*_*g*_ layers that first learns the ligand representation as follows:

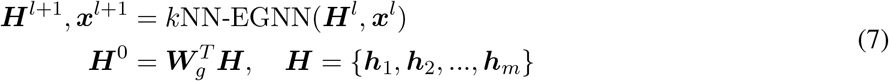

where ***W***_*g*_ ∈ ℝ^*d*×5^ is a learnable mapping matrix. Using the learned representations from both the protein and ligand, the protein-ligand interaction module then predicts the binding probability with a binary cross-entropy loss (**ℒ**_bind_):

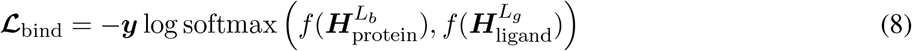

where *f* is a feed-forward neural network that maps the final protein 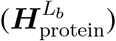 and ligand 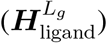 representations to a binding score.

### 4.2 Implementation and Experimental Setup

The protein module of EnzyGen2 comprises three interleaving blocks, each comprising eleven Transformer layers and a single *k*NN-EGNN layer. The Transformer layers were initialized with weights from the pretrained 650M ESM2 model [14]. The protein-ligand interaction module, designed to enable ligand-binding specificity constraint, consists of three *k*NN equivariant graph neural network layers and a two-layer feed-forward neural network. With an embedding dimensionality of *d* = 1, 280 and a nearest neighbor size *k* = 30, the complete architecture consists of 733 million trainable parameters. Our pretraining process was conducted in three distinct stages. First, the model was trained for 1 million steps using a masked language modeling objective, where 20% of residue types and coordinates were masked and subsequently reconstructed. This was followed by an additional 1 million steps of training using only the masked sequence prediction and masked structure reconstruction losses. Finally, the model was further trained for 1 million steps with our full multi-task learning objective, which incorporates the protein-ligand interaction prediction loss. The entire model was trained on eight NVIDIA A6000 GPUs with a batch size of 8,096 tokens. Optimization was performed using the Adam algorithm [81] with a learning rate of 1e-4 and a clip-norm threshold of 1.5. The multi-objective loss function was weighted with hyperparameters *α*=1.0, *β*=0.01, and *λ*=0.5 for the sequence, structure, and binding components, respectively.

For family-specific fine-tuning, the pretrained model was adapted to each enzyme family on a single NVIDIA A6000 GPU for a maximum of 50 epochs, with a learning rate of 5e-5 and a batch size of 1,024 tokens.

During inference, the model generates sequences conditioned on functionally significant residues and NCBI taxonomic identifiers, with ligand information used exclusively during the training phase.

### 4.3 *De novo* Candidate Generation via Nucleus Sampling

Candidate sequences and structures were generated using the best checkpoint from each family-specific fine-tuned model. We employed nucleus sampling [51] (also known as top-*p* sampling) as the decoding strategy. This approach employs a cumulative probability threshold *p* to truncate the predicted amino acid distribution, retaining only the minimal set of amino acids whose collective probability exceeds *p*. By sampling from this restricted distribution, the model maintains high-fidelity sequence generation while avoiding the low-probability “tail” that often leads to non-functional or unstable designs.

For each target enzyme family, we utilized the functionally important residue geometries of all natural homologs as input scaffolds. We generated one *de novo* candidate per input scaffold across a range of threshold values *p* ∈ {0.2, 0.4, 0.6, 0.8}. This multi-threshold sampling strategy was implemented to ensure a diverse set of candidates, traversing different regions of the functional sequence space while adhering to the learned biophysical constraints of the target fold.

### 4.4 Dataset Curation

To develop a robust foundation for ligand-aware sequence-structure co-design, we curated a comprehensive protein-ligand dataset from the PDB and Swiss-Prot databases. These sources were selected because their protein entries are experimentally validated for functionality in laboratory. Specifically, we gathered protein sequences and *C*_*α*_-based backbone structures from PDB, along with sequences from Swiss-Prot and their corresponding structures from the AlphaFold Protein Structure Database. Then we filtered proteins with more than 1,024 amino acids in length, yielding a dataset of 720,993 unique proteins. Binding ligands were then retrieved from UniProtKB [48]. Proteins with an annotated ligand were assigned a positive binding label, while those lacking ligand information were assigned a negative label by randomly sampling a small molecule. Furthermore, we mapped these proteins to 9,002 distinct NCBI taxonomic identifiers retreived from UniProt. For each NCBI identifier, we performed multiple sequence alignments (MSAs) to identify functionally important residues. The dataset was then split into training, validation, and test sets, with 707,843 pairs for training and 6,575 pairs each for validation and testing.

To enable targeted biocatalytic design, we conducted further fine-tuning on three specific enzyme families. To this end, we curated enzyme-specific datasets from the Rhea reaction database by selecting protein entries corresponding to specific chemical reaction IDs: Rhea 18421 for chloramphenicol acetyltransferase (CAT), Rhea 20245 for aminoglycoside adenylyltransferase (AadA) and Rhea 12609 for thiopurine *S*-methyltransferase (TPMT). Protein sequences for each category were retrieved from UniProt [82], and paired with structures from the AlphaFold Protein Structure Database. This resulted in datasets of 3,121 entries for CAT, 2,499 entries for AadA and 6,770 entries for TPMT. To identify functionally important residues, we applied MSA for each category as described earlier. The data for each enzyme family was then randomly divided into training and validation sets in a 9:1 ratio. Fine-tuning was performed exclusively on masked sequence prediction and masked structure reconstruction losses to optimize model performance for each enzyme category.

### 4.5 Automatic Identification of Functionally Important Residues

To facilitate controllable protein design, we define functionally important residues as evolutionarily conserved amino acid patterns within each NCBI taxonomic identifier. This approach leverages the principle that residues critical for structural stability or catalytic activity are subject to stronger selective pressures and thus remain conserved across phylogenetically related sequences.

We used multiple sequence alignment (MSA) to extract these patterns from our curated dataset. Specifically, we employed the ClustalW2 [83] method to perform MSA across all proteins within a given NCBI identifier. A residue at a specific position is designated a functionally important residue if it is present in over 30% of the aligned sequences.

This automated, data-driven strategy bypasses the inherent limitations and lack of scalability associated with manual expert annotation. By identifying evolutionarily preferred amino acid combinations across diverse protein families, this method provides the generative models with precise, site-specific guidance, effectively reducing the combinatorial search space and focusing design efforts on biophysically viable functional folds.

### 4.6 Validation of Designed CAT and AadA

**Cloning**. pET-28a(+) was used as the cloning and expression vector for *de novo* designed CATs and TPMTs in this study. pET-22b(+) was used as the cloning and expression vector for AadA in this study. All the genes were codon-optimized for protein expression in *E. coli* and purchased as plasmids from GeneralBiol.

Functional validation of *de novo* designed antibiotic resistance enzymes. Plasmids carrying the designed antibiotic resistance genes were introduced into *E. coli* BL21(DE3) cells by electroporation individually. After recovery in SOC medium for 45 min at 37 °C, the culture was plated on LB agar containing only the corresponding antibiotic for which the vector confers resistance. For CAT cloned into pET-28a(+), kanamycin was used at a final concentration of 50 *µ*g/mL; for AadA cloned into pET-22b(+), ampicillin was used at 100 *µ*g/mL. Single colonies were picked and cultured overnight in LB medium supplemented with lactose (5 g/L) and the corresponding antibiotic at the specific concentrations (50 *µ*g/mL kanamycin for CAT; 100 *µ*g/mL ampicillin for AadA). After incubation at 37 °C and 250 rpm for 12 h, the resulting cultures were diluted with sterile saline to an OD600 of 1.0 and streaked onto selective LB agar plates supplemented with lactose (10 g/L) and the indicated antibiotic (25 *µ*g/mL chloramphenicol for CAT; 50 *µ*g/mL spectinomycin for AadA). After overnight incubation at 37 °C for 12 h, the activity of the designed antibiotic resistance enzymes was assessed based on the CFUs on the selective plates.

The minimum inhibitory concentration (MIC) of *E. coli* strains expressing the *de novo* designed antibiotic resistance enzymes was determined using a modified broth microdilution assay. Single colonies were inoculated into LB medium supplemented with lactose (5 g/L) and the appropriate selection antibiotic, and cultured at 37 °C with shaking at 250 rpm to prepare the starter culture. For strains expressing CAT, the CAT gene was cloned into the pET-28a(+) vector and maintained in LB medium containing kanamycin (50 *µ*g/mL). For strains expressing AadA, the AadA gene was cloned into the pET-22b(+) vector and maintained in LB medium containing ampicillin (100 *µ*g/mL). Starter cultures were concentrated and adjusted to OD_600_ = 20 to standardize inoculum density and ensure parallelism.

For MIC measurements, 1 *µ*L of the OD_600_ = 20 culture was inoculated into 100 *µ*L LB(lactose) medium containing the antibiotic at a specific concentration in sterile 96-well microtiter plates (final inoculation ratio 1:100). Antibiotic concentrations were optimized to allow for precise determination of inhibitory thresholds.

The plates were incubated at 37 °C for 16 h with shaking at 100 rpm. Bacterial growth was visually evaluated and confirmed by OD_600_ measurement using a Tecan Spark multimode microplate reader. MIC was defined as the lowest antibiotic concentration at which no detectable growth was observed. All assays were conducted in biological triplicate.

For whole-cell acetylation of chloramphenicol by CAT, plasmids encoding previously validated active *de novo* designed variants were introduced into cells by electroporation. The single colonies were picked and cultured overnight in LB medium supplemented with lactose (5 g/L) and kanamycin (50 *µ*g/mL), and cultured at 37 °C with shaking at 250 rpm for 12 hours. The cells were harvested by centrifugation (4,000 x g, 5 min, 4 °C) and were resuspended in M9-N buffer (100 mM, pH 7.4) to an OD_600_ of 0.5. Cell suspension (380 *µ*L) was mixed with chloramphenicol stock solution (20 *µ*L, 50 mg/mL in ethanol) in a sealed vial and incubated at 20 °C with shaking at 200 rpm for 10 min. The reaction was quenched with 400 *µ*L ethyl acetate, vortexed, centrifuged, and the organic phase was collected for UPLC-MS analysis.

UPLC-MS analysis was carried out using a Shimadzu-40D X3 with a 2050 MS detector and an Agilent Poroshell 120 EC-C18 column (4.6 × 50 mm, 4 µm). Water (with 0.1% formic acid) and acetonitrile (MeCN, with 0.1% formic acid) were used as mobile phases.

UPLC-MS analysis method is as follows: an Agilent Poroshell 120 EC-C18 column (4.6 × 50, 4 µm), flow rate = 1.0 mL/min, hold at 5% MeCN (0.1% formic acid) in water (0.1% formic acid) for 0.5 min, ramp up to 95% MeCN (0.1% formic acid) in water (0.1% formic acid) over the course of 7.5 min, hold at 95% MeCN (0.1% formic acid) in water (0.1% formic acid) for 0.5 min, then ramp down to 5% MeCN (0.1% formic acid) in water (0.1% formic acid) over the course of 0.5 min and hold 5% MeCN (0.1% formic acid) in water (0.1% formic acid) for 1 min.

### 4.7 Validation of Designed TPMT

All designed TPMT enzymes described in this study were expressed with an N-terminal 6xHis-tag. A single colony was used to inoculate LB medium containing kanamycin (50 *µ*g/mL). The culture was grown at 37 °C and 250 rpm for 8 h. 3% of this starter culture was used to inoculate the expression culture in TB medium supplemented with kanamycin (50 *µ*g/mL) until OD_600_ reached 0.6 and subsequently induced by the addition of isopropylthio-*β*-D-galactoside (IPTG, final concentration = 0.5 mM). Protein expression was carried out at 20 °C and 200 rpm for 16 h. The cells were finally harvested by centrifugation (4,000 × g, 5 min, 4 °C).

#### Protein solubility analysis

After expression, cells were harvested by centrifugation (4,000 × g, 5 min), resuspended in ice-cold 50 mM NaPi buffer (pH 8.0, 500 mM NaCl), and lysed by sonication using a QSonica Q500 sonicator on ice (2 s on / 4 s off, 45% amplitude, 6 min total). The lysate was clarified by centrifugation (15,000 × g, 20 min, 4 °C) to obtain the soluble fraction (supernatant), while the insoluble pellet was washed once and resuspended in an equal volume of NaPi buffer. Equivalent biomass (normalized by OD_600_) from each fraction and from the whole-cell lysate was mixed with 4 × SDS loading dye containing 10% *β*-mercaptoethanol, heated at 95 °C for 10 min using a Bio-Rad T100 thermal cycler, and then cooled down to room temperature. Denatured protein samples were loaded into the wells of a Bio-Rad Mini-PROTEAN TGX Stain-Free gel (10 *µ*l per well). In total, 2.5 *µ*l PreStained Protein Marker II (10-200 kD) was used as the protein ladder in electrophoresis, and set the voltage on the electrophoresis power supply to a constant voltage of 150 V. Upon the completion of electrophoresis, the SDS-PAGE gel was stained with Coomassie blue by gently microwaving the gel in Bio-Rad Coomassie Brilliant Blue (R-250 Staining Solution purchased from Bio-Rad, catalog number: 1610436) for 30 s. The gel was then destained in a destaining solution (AcOH/MeOH/dd H_2_O=1:2:7 vol/vol/vol) on a microplate shaker (200 rpm) overnight, and analyzed by SDS-PAGE analysis using a Bio-Rad Gel Doc EZ Imager.

#### Protein purification

E. coli cells were resuspended in NaPi buffer (50 mM NaPi buffer, pH 8.0, 500 mM NaCl, 20 mM imidazole). Cells were disrupted by sonication. After pelleting cell debris, the resulting lysate was then centrifuged at 12,000 × g and 4 °C for 60 min using a Lynx 6000 superspeed centrifuge. The supernatant was filtered through a 0.45 *µ*m PES syringe filter and loaded onto a HisTrap HP column (5 mL, GE Healthcare, Piscataway, NJ) using an ÄKTA Start protein purification system (GE Healthcare). Proteins were eluted with an increasing gradient of imidazole from 20 to 500 mM in NaPi buffer at a flow rate of 5.0 mL/min. Fractions containing the desired protein were pooled, and subjected to three exchanges of NaPi buffer (50 mM, pH 8.0) using ultracentrifugal filters (10 kDa molecular weight cut-off, Amicon Ultra, Sigma Millipore) to remove excess salt and imidazole. Concentrated proteins were aliquoted, flash-frozen in liquid nitrogen, and stored at -80 °C with 10 % glycerol as the cryoprotectant until further use. The purity of the protein was further confirmed by SDS-PAGE analysis. Protein concentrations were measured at 280 nm using a NanoDrop OneC spectrophotometer (Thermo Scientific) with buffer as the blank control. Protein concentrations were also further measured using the BCA assay.

#### Michaelis-Menten kinetics measurements

The kinetic measurements were carried out at 25 °C, with each initial rate measured in triplicates. To a 4 mL vial was added 930 *µ*L of NaPi buffer (50 mM, pH 8.0) and 10 *µ*L of the pre-diluted enzyme solution to give a final concentration of 0.526 *µ*M. The mixture was then supplemented with 10 *µ*L of a 100× stock solution of SAH in DMSO and incubated at 25 °C with shaking for 3 min. The reaction was initiated by adding 50 *µ*L of a 200 mM MeI solution in DMSO, mixed briefly, and returned to the shaker (25 °C, 200 rpm). At specified time intervals, 200 *µ*L aliquots were withdrawn and quenched with an equal volume of acetonitrile. After centrifugation (12,000 × g, 5 min), the supernatant was transferred to a 500 *µ*L vial insert, which was then placed in a 2 mL HPLC vial and analyzed by reverse-phase HPLC. Final concentrations of SAH were 2.5, 5, 10, 25, 50, 100, and 250 *µ*M, and the enzyme concentration was 0.5 *µ*M in all reactions.

HPLC Conditions: YMC-Pack Pro C18 column; mobile phase: 10 mM sodium dihydrogen phosphate containing 5 mM sodium 1-heptane sulfonate (pH 3.5) : MeCN=9:1; 1 mL/min; 40 °C; 260 nm. Analytical reverse phase HPLC was conducted using a Shimadzu i-series (66MPa) instructment.

## 5 Data Availability

All datasets used in this study are publicly accessible. Protein sequences and experimentally determined structures were obtained from the PDB (https://www.rcsb.org/docs/programmatic-access/batch-downloads-with-shell-script). Additional protein sequences were retrieved from Swiss-Prot (https://ftp.uniprot.org/pub/databases/uniprot/current_release/knowledgebase/complete/uniprot_sprot.dat.gz), with corresponding structures sourced from the AlphaFold Protein Structure Database (https://alphafold.ebi.ac.uk/download#swissprot-section). Taxonomic identifiers and ligand-binding information were mined from UniProtKB (https://www.uniprot.org/uniprotkb), and ligand structures were downloaded from ChEBI (https://ftp.ebi.ac.uk/pub/databases/chebi/SDF/). For fine-tuning, enzyme-specific data were collected from Rhea, including chloramphenicol acetyltransferase (https://www.rhea-db.org/rhea/18421), aminoglycoside adenylyltransferase (https://www.rhea-db.org/rhea/20245), and thiopurine *S*-methyltransferase (https://www.rhea-db.org/rhea/12609). All datasets were collected up to March 1, 2024. A detailed summary of the curated datasets is provided in Supplementary Table S1, and the curated pretraining and fine-tuning datasets used in this study are available at the data server.

## 6 Code Availability

The code and model checkpoints used in this study are publicly available. The most updated and supported codebase is located at EnzyGen2 codebase.

## Acknowledgment

This work was supported, in part, by an NEC Faculty Research Award (L.L), and National Science Foundation (CHE-2145749 on methyltransferase to Y.Y.; ITE-2448848 on *de novo* protein design to Y.Y.). Y.Y. is an Alfred P. Sloan Research Fellow (FG-2024-22244), a Camille Dreyfus Teacher-Scholar Awardee (TC-25-084), a David & Lucile Packard Fellow (2023-76169) and a Howard Hughes Medical Institute Freeman Hrabowski Scholar.

## Author Contributions

Idealization, Z.S., Y.Z., Y.Y., L.L.; Methodology, Z.S., H.L., Y.Z., Y.Y., L.L.; Data Curation, Z.S.; Computational Experiments, Z.S.; Wet-lab Experiments, H.L.; Software, Z.S.; Analysis of Results, Z.S., Y.Z., Y.Y., L.L.; Manuscript Writing Abstract, Section 1, Z.S., Y.Z; Manuscript Writing Section 2.1-2.5, 3, 4.1-4.5, 5, 6, Z.S.; Manuscript Writing Section 2.6, Section 4.6, 4.7, H.L., Y.Z.; Candidate Selection for AadA, CAT, Y.Z.; Candidate Selection for TPMT, H.L; Editing, Z.S., H.L., Y.Z., Y.Y., L.L.; Funding Acquisition, L.L., Y.Y.

