## Supplemental material for "Co-designing sequence and structure of functional *de novo* enzymes with EnzyGen2"

### (SUPPLEMENTAL INFORMATION)

| Database | Data Size | Purpose for Model Training |
|---|---|---|
| Protein Data Bank (PDB) | 202,433 | Using the sequence and $C_\alpha$ backbone structures for model learning, which provides the supervised data for sequence-structure co-design |
| Swiss-Prot | 542,378 | Experimentally validated protein sequences used for model learning |
| AlphaFold Structure Database | 542,378 | AlphaFold2 predicted structures for proteins from Swiss-Prot to achieve model training |
| UniprotKB | 720,993 | Collecting the binding ligand information and the NCBI taxonomy identifier for each protein |
| ChEBI | 153,139 | Extracting ligand structures and features used for protein-ligand interaction constraint |
| Rhea | 3,121; 2,499; 6,770 | Collecting fine-tuning data for the enzyme families CAT, AadA and TPMT |

**Table S1:** EnzyGen2 pretraining and fine-tuning datasets curated from a list of publicly available data sources.

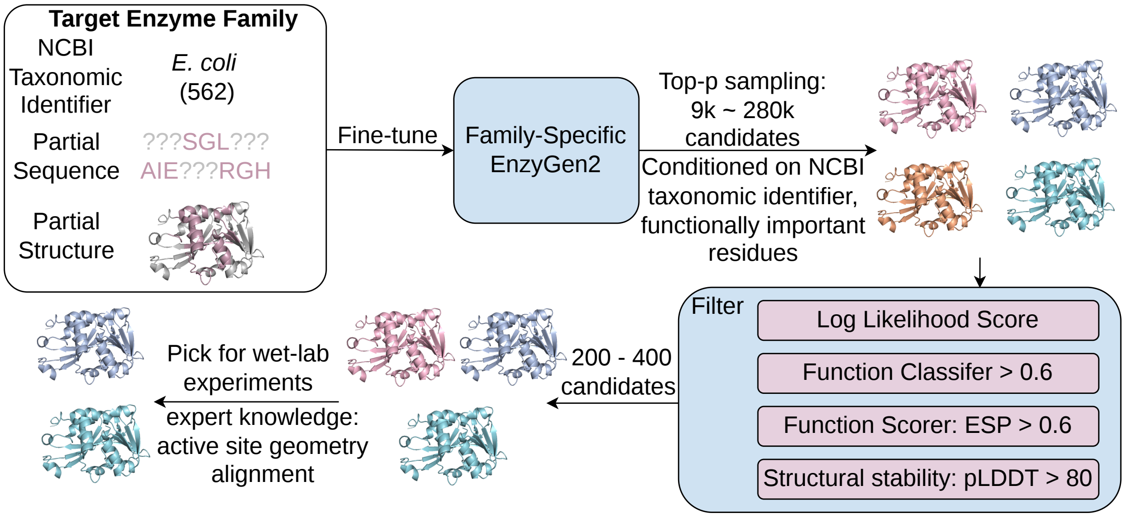


**Fig. S1 Systematic screening of *de novo* enzyme candidates.** The selection workflow employs a multi-stage approach to identify functional leads for experimental validation. Initially, the pretrained EnzyGen2 is fine-tuned on a specific enzyme family targeted for subsequent wet-lab testing. Candidate sequences are then generated using nucleus sampling. These candidates are then filtered based on a predefined suite of scoring metrics. From the refined set of candidates, we make final selections for wet-lab validation, prioritizing sequences with either high predictive confidence or high sequence novelty relative to natural homologs.

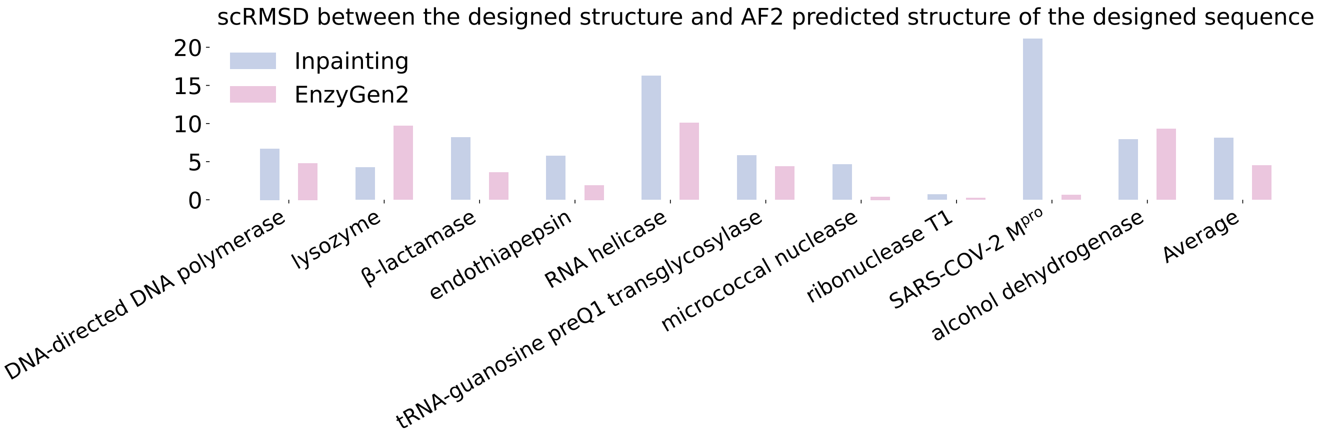


**Fig. S2 Self consistency between designed structures and sequences.** We evaluate the self-consistency (scRMSD) of our model by comparing the RMSD between the designed structures and the structures predicted by AlphaFold2 from the corresponding designed sequences. Results are shown for the ten largest enzymes in the test set, enabling a robust comparison between the simultaneous sequence-structure co-design baseline (Inpainting) and EnzyGen2.

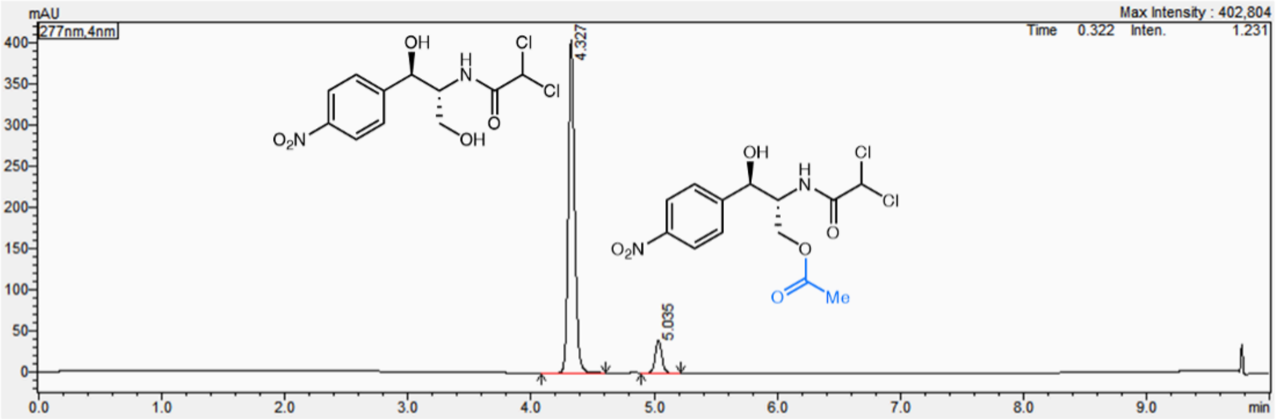


| Peak# | Ret. Time | Area% |
|---|---|---|
| 1 | 4.327 | 90.766 |
| 2 | 5.035 | 9.234 |
| Total | | 100.000 |

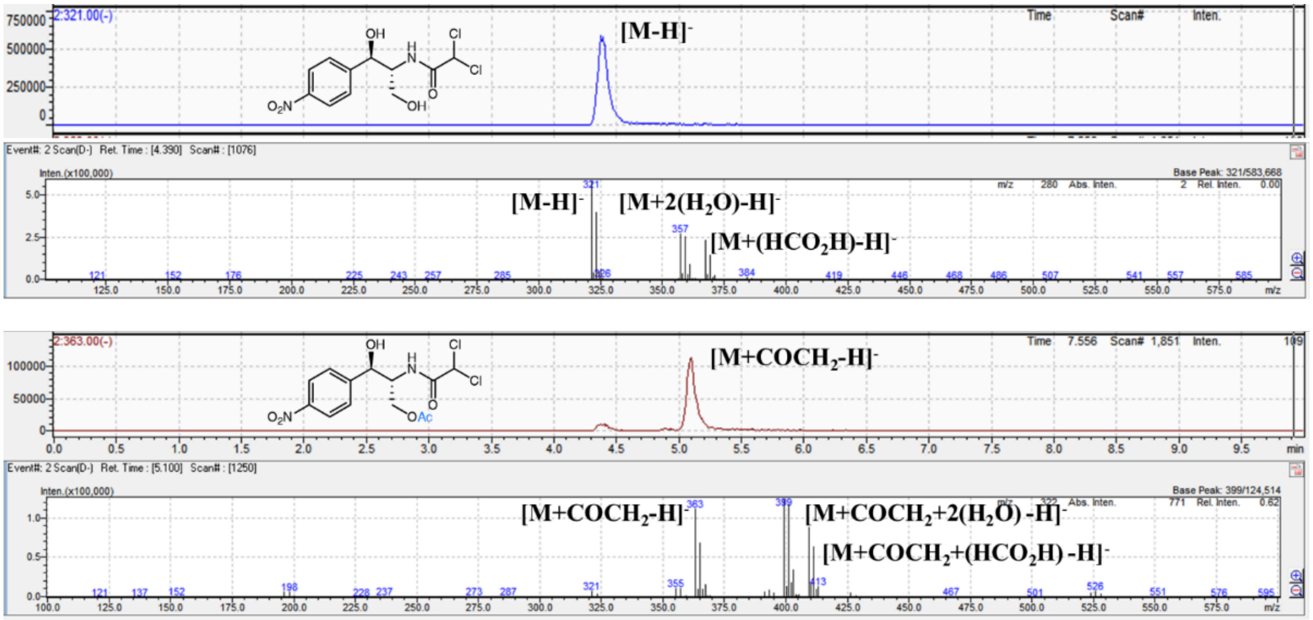


**Fig. S3 UPLC-MS trace of whole-cell acetylation reaction of chloramphenicol.** The trace of wild-type CAT.

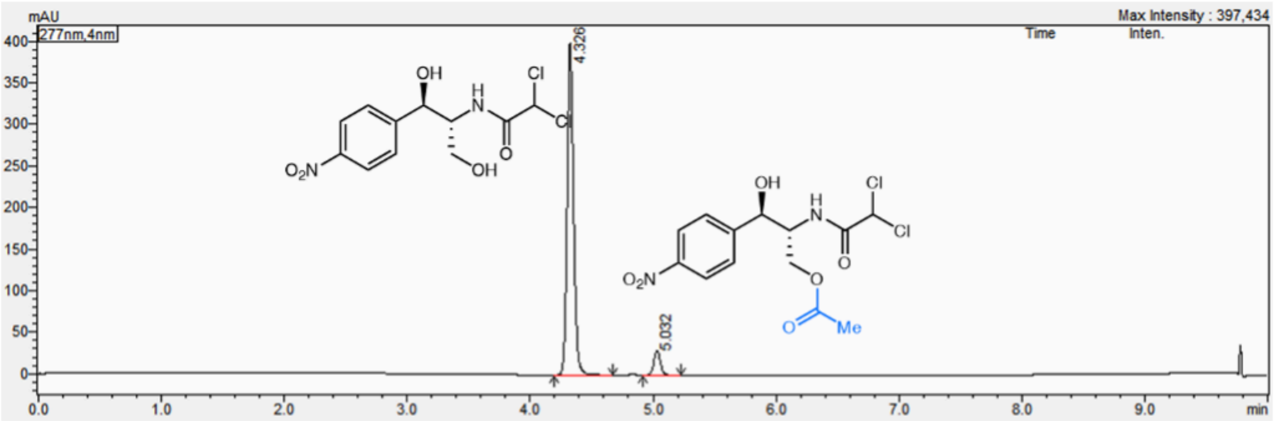


| Peak# | Ret. Time | Area% |
|---|---|---|
| 1 | 4.326 | 92.860 |
| 2 | 5.032 | 7.140 |
| Total | | 100.000 |

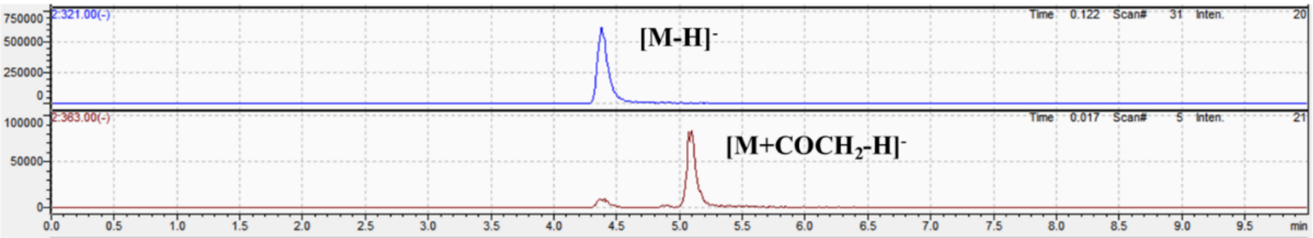


**Fig. S4 UPLC-MS trace of whole-cell acetylation reaction of chloramphenicol.** The trace of CAT-8.

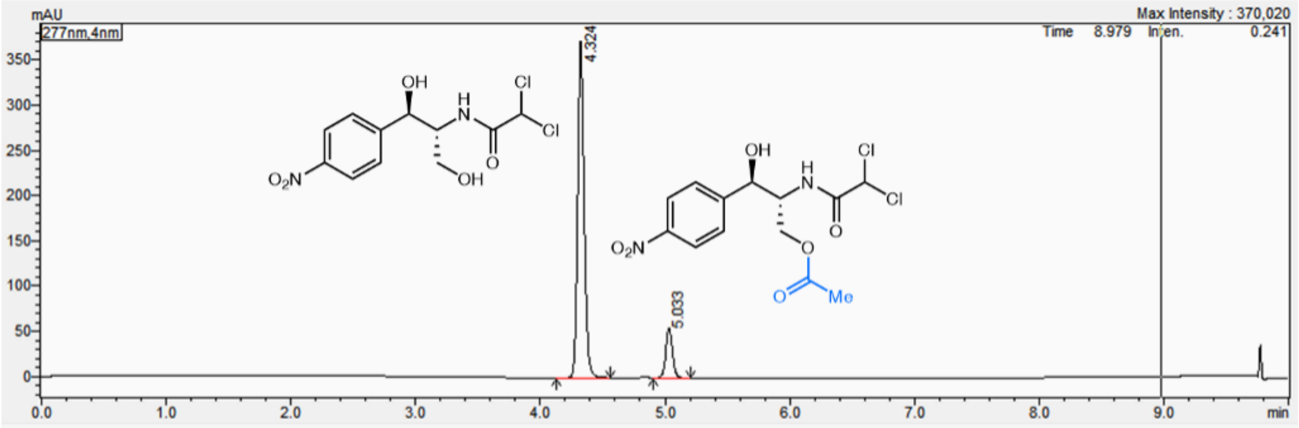


| Peak# | Ret. Time | Area% |
|---|---|---|
| 1 | 4.324 | 86.711 |
| 2 | 5.033 | 13.289 |
| Total | | 100.000 |

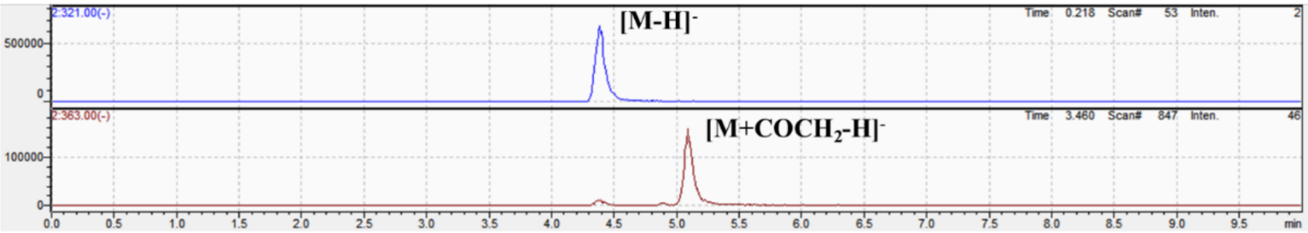


**Fig. S5 UPLC-MS trace of whole-cell acetylation reaction of chloramphenicol.** The trace of CAT-11.

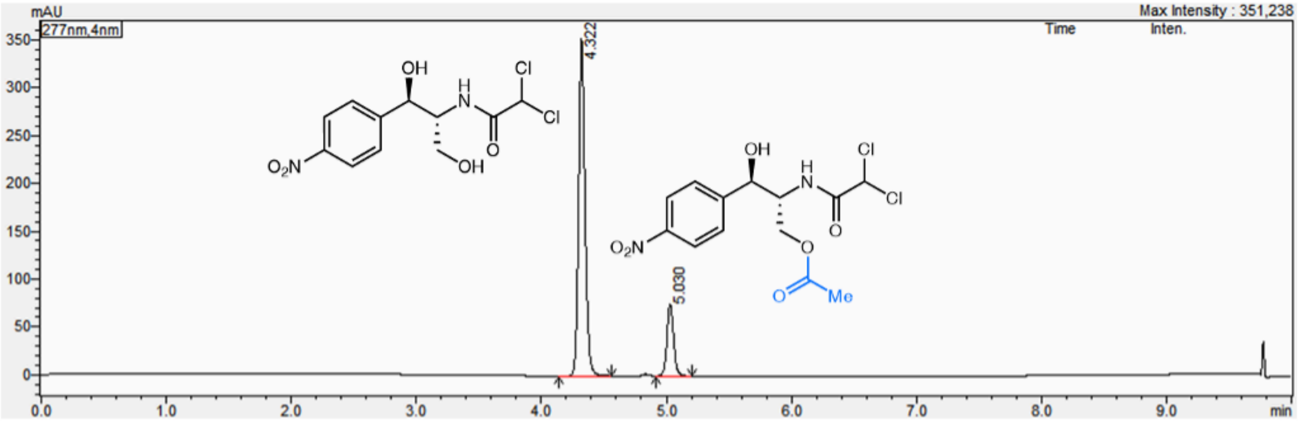


| Peak# | Ret. Time | Area% |
|---|---|---|
| 1 | 4.322 | 82.078 |
| 2 | 5.030 | 17.922 |
| Total | | 100.000 |

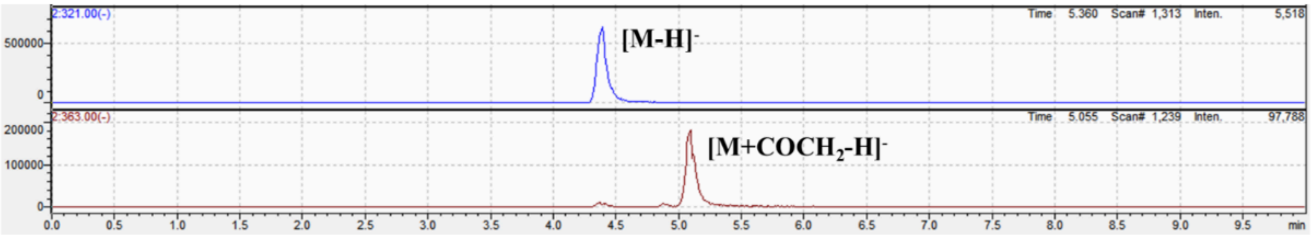


**Fig. S6 UPLC-MS trace of whole-cell acetylation reaction of chloramphenicol.** The trace of CAT-16.

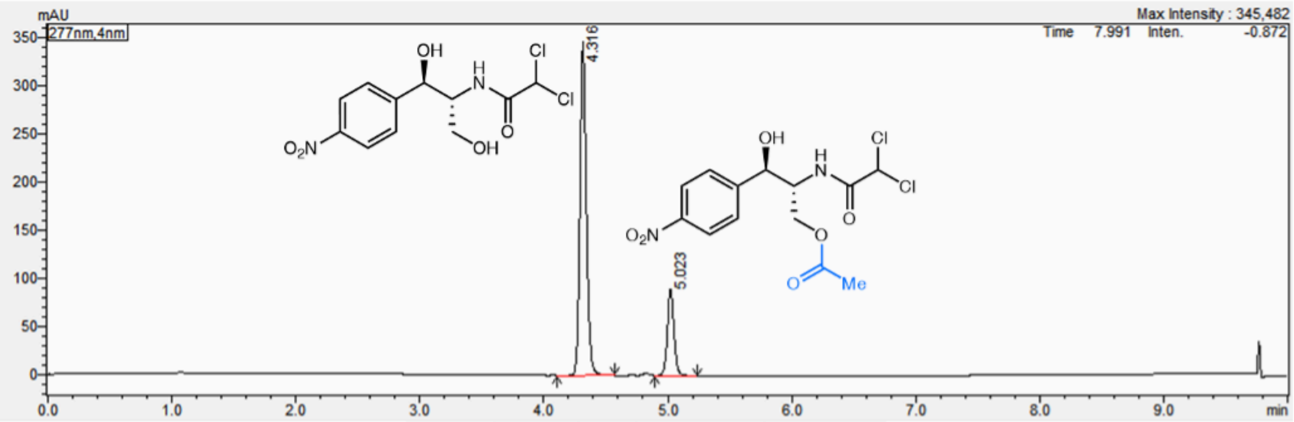


| Peak# | Ret. Time | Area% |
|---|---|---|
| 1 | 4.316 | 79.025 |
| 2 | 5.023 | 20.975 |
| Total | | 100.000 |

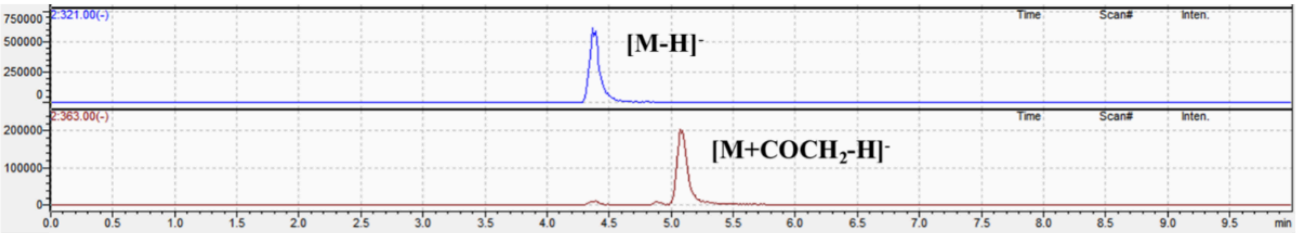


**Fig. S7 UPLC-MS trace of whole-cell acetylation reaction of chloramphenicol.** The trace of CAT-17.

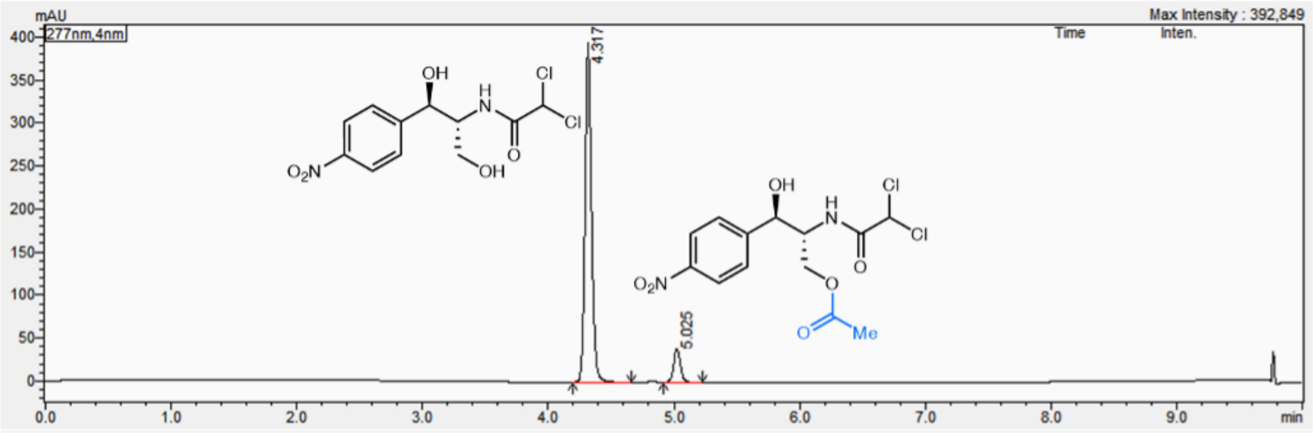


| Peak# | Ret. Time | Area% |
|---|---|---|
| 1 | 4.317 | 90.669 |
| 2 | 5.025 | 9.331 |
| Total | | 100.000 |

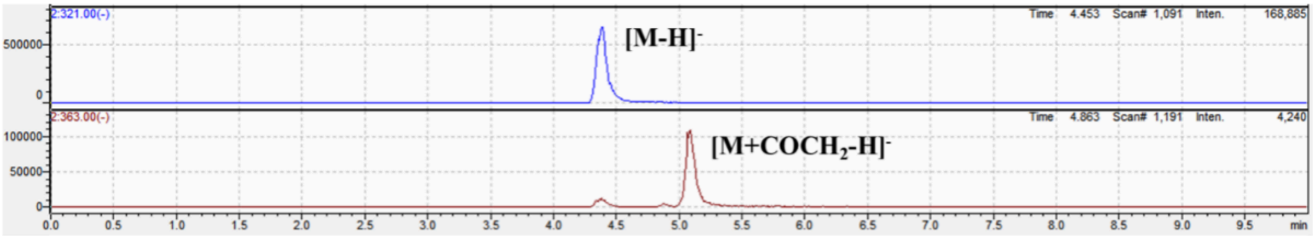


**Fig. S8 UPLC-MS trace of whole-cell acetylation reaction of chloramphenicol.** The trace of CAT-19.

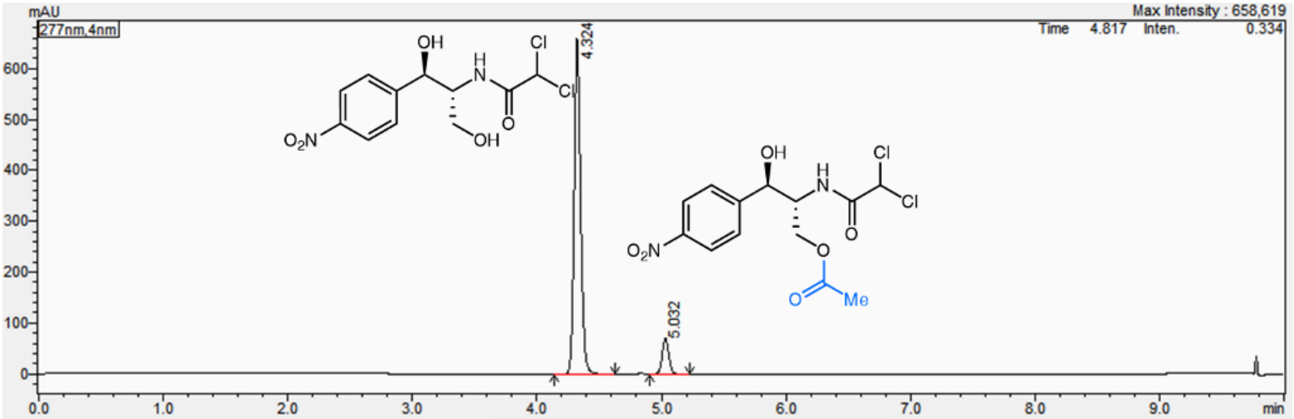


| Peak# | Ret. Time | Area% |
|---|---|---|
| 1 | 4.324 | 90.040 |
| 2 | 5.032 | 9.960 |
| Total | | 100.000 |

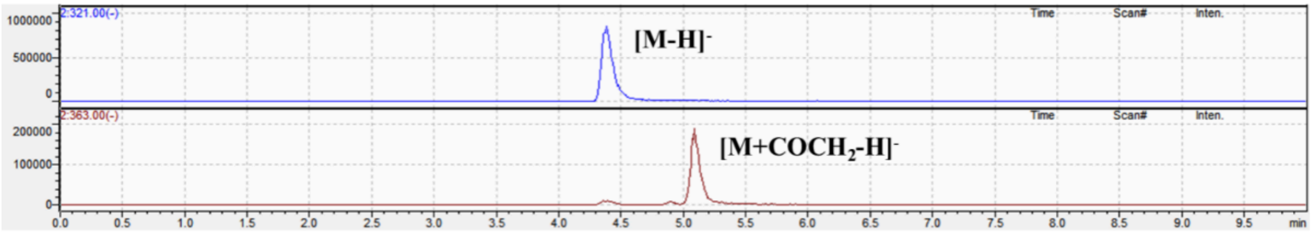


**Fig. S9 UPLC-MS trace of whole-cell acetylation reaction of chloramphenicol.** The trace of CAT-20.

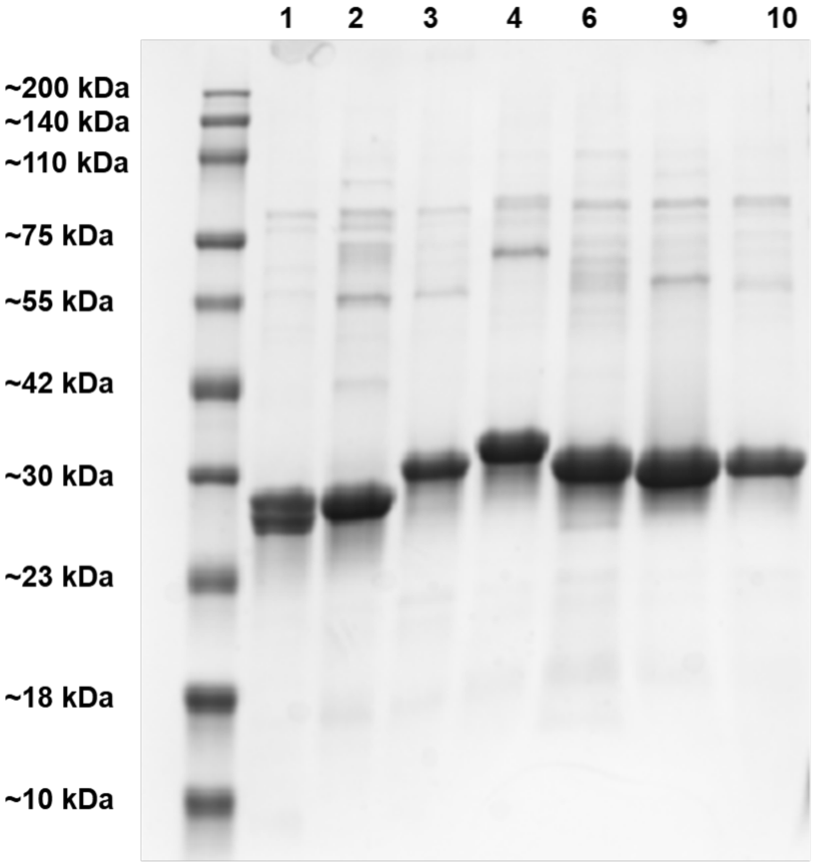


**Fig. S10 SDS-PAGE of purified *de novo* designed TMPTs.** From left to right, the figure shows the bands of purified TMPT-1, TPMT-2, TPMT-3, TPMT-4, TPMT-6, TPMT-9 and TPMT-10. The presence of a lower-molecular-weight band beneath the primary band of TPMT-1 indicates possible protein degradation or premature translational termination.
